# The lncRNA PACER Regulates Lung Adenocarcinoma Phenotypes via COX-2 Signaling and RNA Structural Dynamics

**DOI:** 10.64898/2025.12.17.694931

**Authors:** Samuel Z. Desind, Samira K. Bell, Robert L. Merritt, Benjamin D. Carr, Hannah K. Shorrock, Carol S. Lutz

**Affiliations:** Department of Microbiology, Biochemistry, and Molecular Genetics, Rutgers Biomedical & Health Sciences, New Jersey Medical School, School of Graduate Studies, Newark, NJ 07103; The RNA Institute, College of Arts & Sciences, University at Albany, State University of New York, University at Albany, Albany, NY 12222; Department of Biological Sciences, College of Arts & Sciences, University at Albany, State University of New York, University at Albany, Albany, NY 12222

**Keywords:** PACER, COX-2, LncRNA, Lung adenocarcinoma, SHAPE-MaP

## Abstract

Long noncoding RNAs (lncRNAs) are a class of regulatory RNAs with critical roles in cellular homeostasis and disease pathogenesis, including cancer. Dysregulated lncRNA expression in lung adenocarcinoma (LUAD) contributes to oncogenic mechanisms through immune and inflammatory signaling networks, including the arachidonic acid (AA) signaling pathway. Here, we investigate the functional significance of the lncRNA PACER (*PTGS2* Antisense NF-κB1 Complex-Mediated Expression Regulator), which modulates cyclooxygenase-2 (COX-2), a key enzyme in the AA pathway.

Using stable shRNA-mediated knockdown in A549 LUAD cells, we show that PACER silencing reduces COX-2 expression and impairs cellular proliferation, migration, and invasion. Bioinformatic analysis using LncLOOM revealed conserved motifs across primates and mice, including predicted binding sites for miR-18a-5p, miR-196-5p, miR-1306-5p, and miR-423-3p. To define PACER’s structural organization, we performed selective 2′-hydroxyl acylation analyzed by primer extension and mutational profiling (SHAPE-MaP), generating the first secondary structure determination of PACER. SHAPE-MaP and SuperFold analyses revealed a compact, stabilized 5′ domain and a flexible central and 3′ region. ΔSHAPE analysis identified multiple sites of differential protection, and Rsample clustering resolved two dominant in vivo conformations, suggesting that PACER adopts a modular architecture with defined structural domains. These structured regions coincide with conserved motifs associated with NF-κB p50 and predicted miRNA binding sites, indicating that PACER function may depend on conformational switching that modulates protein and miRNA accessibility. Together, these findings establish PACER as a regulator of LUAD proliferation and invasion via COX-2 signaling and highlight its potential as a biomarker and therapeutic target in inflammatory cancers.

## Introduction & Background

Globally, lung and bronchial cancers are the leading cause of cancer mortality, accounting for 18.7% of cancer deaths and 12.4% of new cancer cases (Bray et al. 2024). In the United States, there were an estimated 234,580 new cases and 125,070 deaths from lung and bronchus cancer in 2024 (*American Cancer Society. Cancer Facts & Figures* 2025). Non-small cell lung cancer (NSCLC) comprises ∼85% of total lung cancer cases, with lung adenocarcinoma (LUAD) and lung squamous cell carcinoma (LUSC) as the predominant subtypes (Ryan 2018). LUAD is commonly associated with chronic inflammation, and the majority of identified LUAD tumors are invasive, driven by common EGFR mutations, ALK rearrangements, and other common oncogenic markers (Molina et al. 2008; Myers & Wallen 2020; Travis et al. 2013). Patient genetic background and the presence of treatment-resistant mutations can significantly impact the efficacy of traditional cancer treatment strategies, targeted chemotherapies and immunotherapy (E. S. Kim 2016; Ko et al. 2017; Tsvetkova & Goss 2012; Wu et al. 2020).

One inflammatory signaling pathway commonly dysregulated in lung cancers is the arachidonic acid (AA) pathway (Figure 1), which synthesizes a class of molecules called eicosanoids. Cyclooxygenase 2 (COX-2, gene name *PTGS2*), a critical enzyme within the AA pathway, plays an essential regulatory role in this inflammatory response and exhibits overexpression across various cancers, including breast, ovarian, colorectal, and lung cancers (Ferrandina et al. 2002; Half et al. 2002; Lutz & Cornett 2013; Petkova et al. 2004; Roelofs et al. 2014). COX-2 converts AA into prostaglandin H2 (PGH_2_) and acts as the rate-limiting enzyme in the production of prostaglandin E2 (PGE_2_) (Hanna & Hafez 2018). PGE_2_ binds to one of four distinct G protein-coupled receptors, triggering cAMP/PKA signaling cascades regulating multiple downstream pathways, including MAPK/Erk, PI3K, AKT, and β-catenin signaling (Castellone et al. 2005; Dalwadi et al. 2005; Krysan et al. 2005). In cancer, chronic activation of these pathways results in pro-tumorigenic processes, including increased cell proliferation and migration, angiogenesis, and cancer cell immune evasion (Finetti et al. 2020; Hangai et al. 2016) (Figure 1). Regulation of COX-2 expression is tightly controlled at multiple levels, including promoter and enhancer elements containing transcription factor binding sites, microRNA (miRNA)-mediated repression through interactions with the 3′ UTR, alternative polyadenylation, control by long noncoding RNAs (lncRNAs) and regulation of overall RNA stability (Desind et al. 2023; Dixon 2003; FitzGerald 2003; Lutz & Cornett 2013; Monteleone & Lutz 2020). An important enhancer of cyclooxygenase-2 (COX-2) is the lncRNA *PTGS2* antisense NF-κB1 complex-mediated expression regulator RNA known as PACER (f.k.a. the p50-Associated COX-2 Extragenic RNA) (Figure 1).

**Figure 1.**
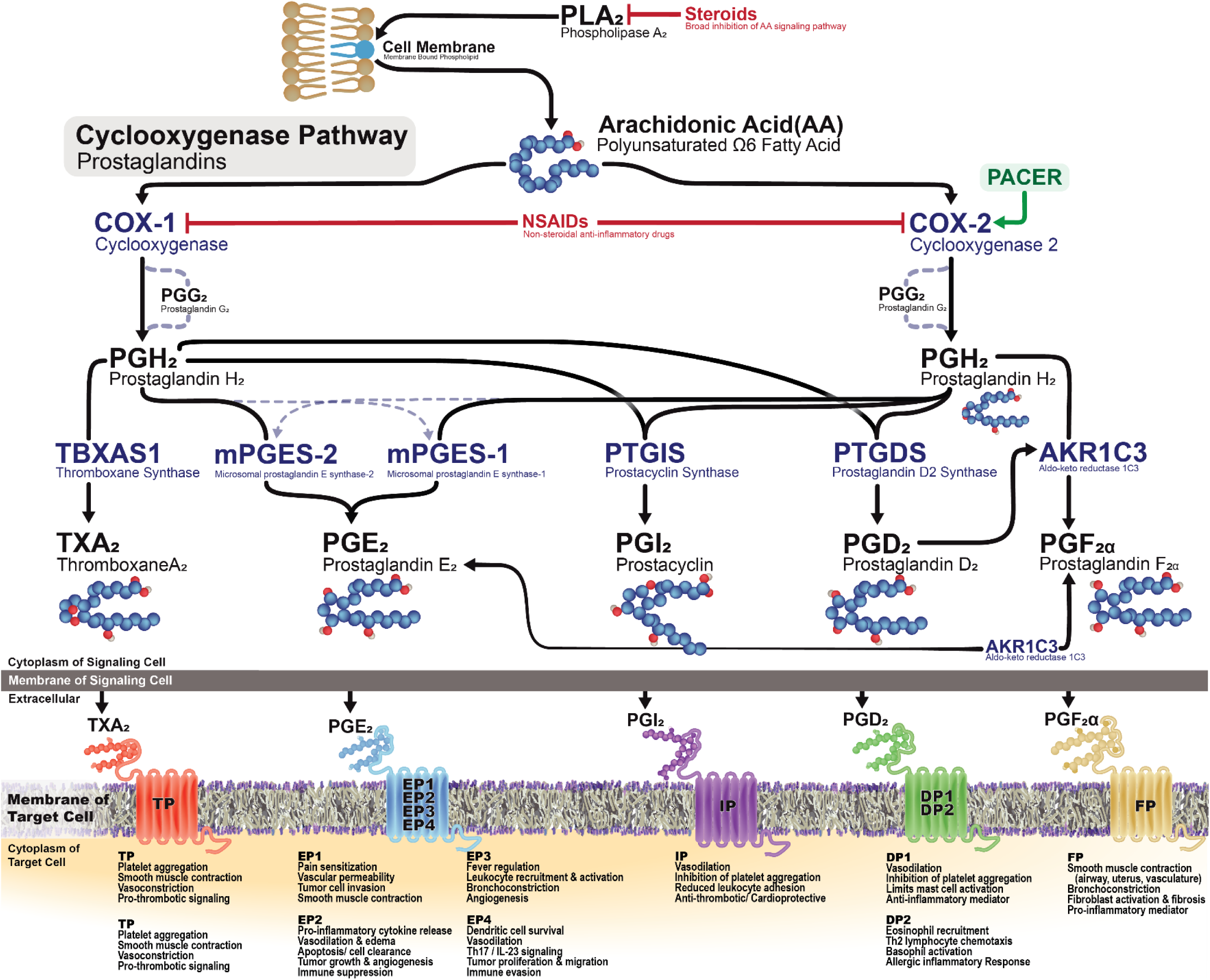
Cyclooxygenase Pathway & Prostaglandin Receptor Signaling. Arachidonic acid (AA), released from membrane phospholipids by phospholipase A₂ (PLA₂), is converted by cyclooxygenase-1 and -2 (COX-1/COX-2) into the intermediate PGH₂, which is further metabolized by specific synthases into thromboxane A₂ (TXA₂), prostaglandin E₂ (PGE₂), prostacyclin (PGI₂), prostaglandin D₂ (PGD₂), and prostaglandin F₂α (PGF₂α). These lipid mediators act on distinct G-protein-coupled receptors in target cells (TP, EP1-EP4, IP, DP1/DP2, FP) to regulate vascular tone, platelet function, leukocyte recruitment, and inflammatory responses. Steroids inhibit AA release at PLA₂, while non-steroidal anti-inflammatory drugs (NSAIDs) block COX activity. The long noncoding RNA PACER (green) selectively promotes COX-2 expression, linking transcriptional regulation to prostaglandin production. The downstream regulatory roles of cyclooxygenase pathway metabolites are highly cell-type and context-dependent, contributing to both homeostatic and chronic inflammation. **Abbreviations:** TP, thromboxane receptor; EP1-EP4, PGE₂ receptors 1-4; IP, prostacyclin receptor; DP1/DP2, PGD₂ receptors 1/2; FP, PGF₂α receptor

LncRNAs are defined as RNAs with a minimum length of 200 nucleotides that do not code for proteins; however, this broad class of RNAs displays extensive structural, functional, and mechanistic diversity. LncRNAs regulate the gene expression of intrinsic and extrinsic cell signaling pathways, influencing growth, motility, immune signaling, apoptosis and many other aspects of cellular metabolism. The abundance of lncRNAs in the human genome has inevitably led to their discovery in virtually every hallmark of cancer, solidifying their significance in the complex regulation of tumorigenesis (Gutschner & Diederichs 2012; Huarte 2015; J. Wang et al. 2020). Typically, lncRNAs are transcribed from the promoter regions, intronic sequences or antisense sequences of protein-coding genes by RNA polymerase II (Kornienko et al. 2013; Mattick et al. 2023). Indeed, the lncRNA PACER is positioned on chromosome 1, lying antisense and overlapping the 3′ promoter region of the COX-2 gene.

Advancements in the sensitivity and selectivity of experimental and bioinformatic techniques have increased the rate of discovery of many novel lncRNAs, while improved sequencing strategies and structural modeling techniques have facilitated the characterization of distinct lncRNA motifs and their physiological importance (Guo & Guttman 2022; Huang et al. 2011; Ross et al. 2021; Smola & Weeks 2018). As part of this improved understanding of lncRNAs, selective 2′-hydroxyl acylation analyzed by primer extension and mutational profiling (SHAPE-MaP) has been widely used to analyze their structures, including MALAT1, MEG3, and many others (Manfredonia et al. 2020; Monroy-Eklund et al. 2023; Sherpa et al. 2018; Siegfried et al. 2014).

Krawczyk and Emerson were the first to identify and characterize the lncRNA PACER in humans (Krawczyk & Emerson 2014). Chromatin immunoprecipitation analysis revealed robust RNA polymerase II (RNAP II) binding upstream of the COX-2 transcriptional start site, corresponding to the PACER loci. They also identified CTCF/cohesin binding sites around the COX-2 locus. The CTCF/cohesin interactions are thought to maintain the structural stability of these loci, granting RNA polymerase II efficient access to both the PACER and COX-2 promoter regions. RNA binding and ChIP-PCR assays showed that PACER regulates COX-2 mRNA by interacting with the NF-κB p50 subunit, which constitutively binds the COX-2 promoter in a transcriptionally repressive manner. Finally, they demonstrated that lipopolysaccharide (LPS)-induced immune stimulation increased PACER transcription, sequestration of inhibitory NF-κB p50 and binding of the p50/p65 NF-κB heterodimer to the COX-2 promoter, enhancing COX-2 transcription (Krawczyk & Emerson 2014). While the authors provided convincing evidence that PACER interacts with p50, they did not discuss or identify where or how this interaction occurs.

Our previous work demonstrated that PACER is upregulated in lung adenocarcinoma cells (LUAD), showing a strong positive correlation with COX-2 expression and an association with decreased disease-specific survival in high-risk patient subgroups (Desind et al. 2022). A series of pharmacologic experiments in A549 cells further revealed a feedback loop within the PACER/COX-2/PGE₂ axis. Celecoxib suppressed PACER expression by approximately 75% (t(2.55) = 10.71, p = 0.003), while PGE₂ significantly increased both PACER and COX-2 expression (t(3.784) = 4.120, p = 0.016, COX-2 (t(3.99) = 6.34, p = 0.003). Treatment with TXA₂ had no significant effect on PACER or COX-2 expression (p > 0.05), indicating the specificity of this regulatory circuit towards PGE_2_ (Desind et al. 2022). Together, this research suggests that dysregulation of PACER may contribute to the overall disease phenotype across multiple cancer types, including LUAD (Desind et al. 2022; Qian et al. 2016; Sun et al. 2021).

In this study, we sought to better understand the role of PACER in shaping the overall disease phenotype in LUAD. To investigate the role of PACER in tumorigenic phenotypes, we generated stable PACER knockdown cell lines and employed functional assays of proliferation, migration, and invasion. We used LncLOOM to assess evolutionary conservation of short functional motifs, and SHAPE-MaP to probe PACER structure, comparing in vivo and ex vivo conditions to identify regions of structural importance that may contribute to PACER’s regulatory activity. This study combines functional assays, bioinformatic models of conservation, and chemical probing experiments to demonstrate the role of PACER in lung cancer tumorigenesis and to identify structural features relevant to PACER’s function.

## Results

### ShRNA-Mediated PACER Knockdown Impairs NF-κB/COX-2 Signaling and Proliferation

To assess the functional role of PACER in LUAD, we sought to knockdown PACER expression in A549 cells, a human lung adenocarcinoma cell line that robustly expresses COX-2 and PACER (Cornett & Lutz 2014; Desind et al. 2022). Using short hairpin RNA (shRNA) targeting PACER (shPACER), we generated A549 cell lines with constitutive knockdown of PACER expression through lentiviral transduction (Sup. Figure 1, Figure 2A). Alongside the knockdown cell lines, experiments were performed with wild-type cells and cells transduced with a non-mammalian control shRNA (shNMC, Table 1). The non-mammalian control shRNA does not target any known transcripts in mammalian cells. Transduced cells were selected using puromycin (5 µg/mL).

**Figure 2.**
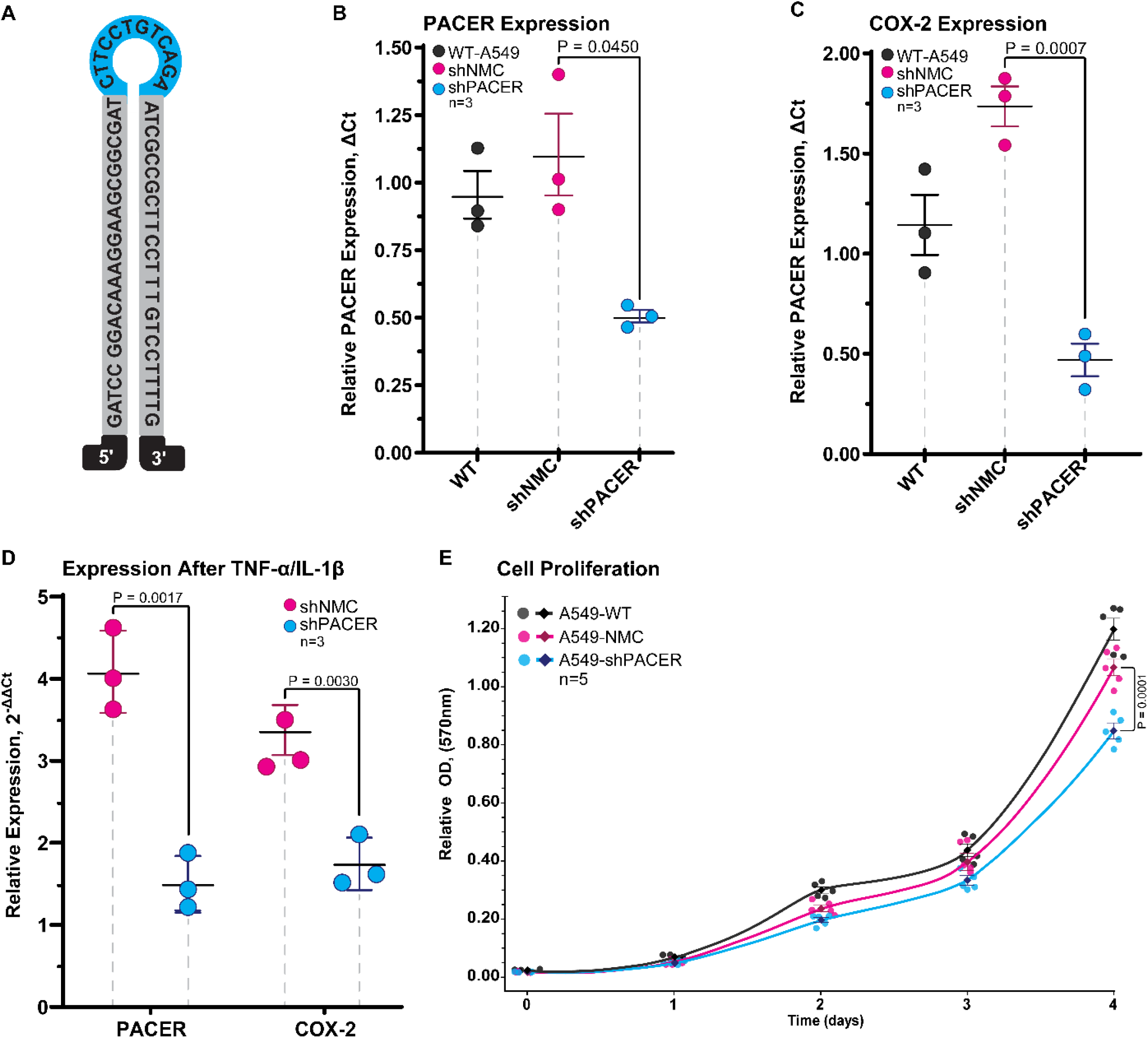
Stable shRNA Knockdown of PACER in A549 Cells Impacts COX-2 Expression & Cell Proliferation. **(A)** Schematic representation of PACER shRNA 1. **(B)** Quantitative RT-PCR analysis of PACER expression in wild-type (WT), non-mammalian shRNA control (shNMC), and PACER knockdown (shPACER) A549 cells. PACER expression was significantly reduced in KD cells versus shNMC (t(2.10, n=3) = 4.34, p = 0.0450). **(C)** COX-2 expression levels in the same samples under basal conditions demonstrate significantly reduced COX-2 expression in shPACER cells (t(3.84, n=3) = 9.86, p = 0.0007). Baseline expression was normalized to GAPDH. **(D)** Pro-inflammatory cytokine treatment (50 ng/mL TNF-α and 10 ng/mL IL-1β in serum-free medium) increased PACER and COX-2 expression in WT and shNMC cells, while shPACER cells had a significantly dampened expression (PACER; t(2.44, n=3) = 12.05, p = 0.0031, COX-2; t(3.30, n=3) = 9.02, p = 0.0020). Data were normalized to GAPDH expression and relative to untreated controls. Plots show individual data points, the mean (center black bar), and error bars, representing SEM. **(E)** Cell proliferation was measured by optical density (OD) after MTT treatment. Cells were seeded at 3.0×104 cells per well in a 24-well plate. MTT was added at a final concentration of 0.5 mg/mL. Optical density (OD) was measured at a wavelength of 570 nm, and the background was subtracted. At day 4, the shPACER cells exhibited a significantly lower OD compared to both WT (t(6.53, n=5) = 8.13, p = 0.0001) and NC (t(7.76, n=5) = 7.16, p = 0.0001). P-values were calculated using a two-tailed Welch’s t-test (α = 0.05).

**Table 1.**
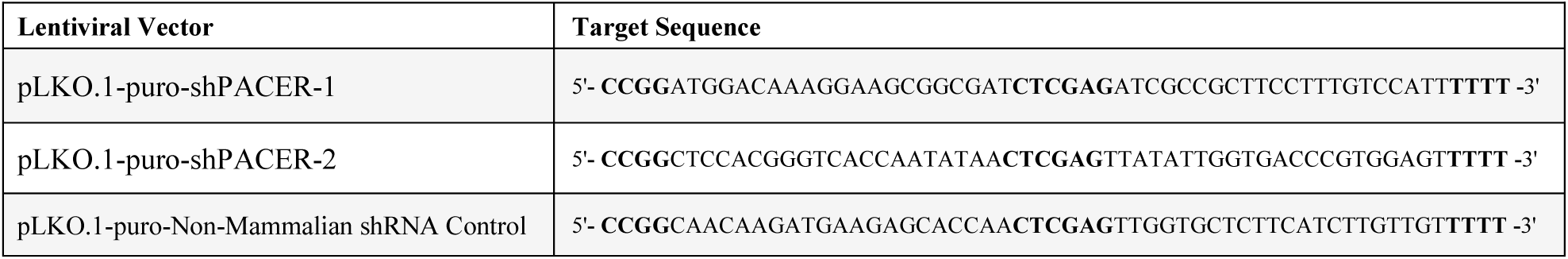
Lentiviral Vectors for shRNA PACER Knockdown & Non-targeting Controls. These RNAi vectors were used to generate lentiviral transduction particles carrying either an shRNA sequence targeting the PACER gene or a non-mammalian control (NMC) sequence. The shRNA in the pLKO.1-puro-shPACER-1 is positioned 601 nt downstream of the PACER transcriptional start site as described by previous studies (Krawczyk & Emerson, 2014; Qian et al., 2016). The pLKO.1-puro-shPACER-2 begins 638 nt from the start of PACER. The 5’ flank sequence (CCGG), loop sequence (CTCGAG), and 3’ flank sequence (TTTT) are indicated in bold. The pLKO.1-puro-Non-Mammalian shRNA Control contains a target sequence that does not match any known human RNA sequence.

PACER transcript levels were measured using quantitative real-time PCR (qRT-PCR), which confirmed that shRNA transduction resulted in significantly reduced PACER RNA expression relative to both WT and shNMC controls (t(2.10, n=3) = 4.34, p = 0.0450) (Figure 2B). ShRNA knockdown achieved greater than 50% knockdown efficiency relative to shNMC expression. As expected, PACER expression did not significantly differ between shNMC and WT cells, indicating the knockdown effects of the PACER shRNA construct are a result of the shRNA specificity for PACER. Consistent with our earlier siRNA-based findings (Desind et al. 2022), PACER knockdown led to a significant decrease in COX-2 transcript expression (t(3.84, n=3) = 9.86, p = 0.0007), with a reduction of approximately two-thirds compared to control cell lines (Figure 2C).

We next investigated the effects of pro-inflammatory cytokine-mediated NF-κB upregulation in shPACER knockdown cells. Cells were starved and cultured overnight. The following day, 50 ng/mL TNF-α and 10 ng/mL IL-1β were added. After 12 hours of treatment, we measured PACER and COX-2 transcript levels by qRT-PCR. We found both PACER and COX-2 to be upregulated in WT and shNMC cells compared to untreated controls, and shPACER cells showed a significantly reduced response, with decreased expression of both PACER and COX-2 (t(2.44, n=3) = 12.05, p = 0.0031, t(3.30, n=3) = 9.02, p = 0.0020) compared to shNMC controls (Figure 2D). This suggests that PACER knockdown effectively reduces COX-2 transcription in a pro-inflammatory environment induced by cytokine stimulation.

Sustained proliferative signaling is a hallmark of cancer and a key driver of tumor growth and progression in lung cancer (Hanahan 2022; Hanahan & Weinberg 2011; Park & Hong 2016). Given this central role in tumor development, we investigated whether PACER knockdown affects the proliferation of lung cancer cells. We measured the relative optical density (OD) of A549-WT, shNMC, and shPACER cells after methyl thiazolyl tetrazolium (MTT) treatment over four days. The shPACER knockdown cell line had consistently lower relative OD compared to both A549-WT and shNMC control cells, indicating a reduction in metabolic activity and cell proliferation (Figure 2E). Both WT and shNMC cultures exhibited increased OD through Day 4, while we observed reduced proliferation in shPACER cells beginning at Day 2. By day 4, the shPACER cells exhibited a significantly lower OD compared to both WT (t(6.53, n = 5) = 8.13, p = 0.0001) and shNMC (t(7.76, n = 5) = 7.16, p = 0.0001). This ∼24% decrease compared with shNMC cells demonstrates that PACER knockdown reduced the proliferative capacity of lung cancer cells.

### PACER knockdown Suppresses Migration and Invasion of Lung Cancer Cells

To further investigate the impact of PACER knockdown on the lung cancer cell phenotype, we performed wound-healing or migration assays. We used wound healing assays to measure cells’ ability to migrate after the formation of a “wound” or clearing. WT-A549, shNMC and shPACER cells were plated in 24-well plates and grown to near confluency. A wound was created by scoring the surface with a 200 µL micropipette tip, followed by several washes to remove cells and debris from the wound (n = 6). We imaged cell migration into the cleared area at 0, 12, 24, and 48 hours post-wound formation. The change in growth area was measured relative to the 0-24-hour time interval.

Figure 3A shows the original image and area quantification analysis for the 0- and 24-hour time points as a reference. Imaging and analysis of the 0-24 hour time interval showed a significantly reduced migration of shPACER cells relative to shNMC controls (t(10, n = 6) = 2.338, p = 0.0415) (Figure 3B). By 24 hours, both A549 WT and shNMC cells had covered 61 and 55% of the wound area, respectively, versus 43% in the shPACER cells, relative to the initial wound size (Figure 3C). ShPACER cells exhibited visibly reduced coverage, especially against the shNMC cells, indicating PACER knockdown significantly impacted cell migration.

**Figure 3.**
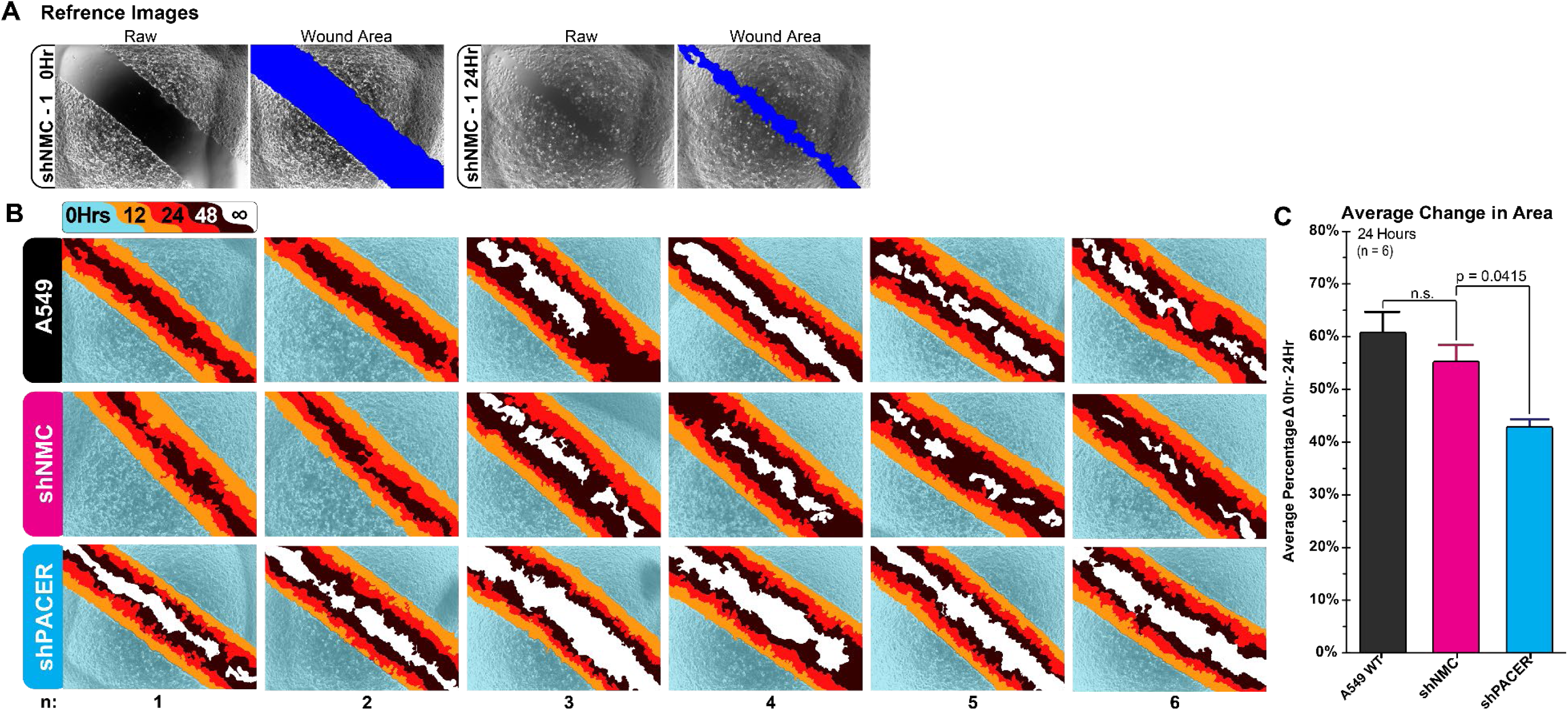
ShPACER KD Wound Healing Assays. A549 cells were grown in a 24-well plate and scored using a 200 μL pipette tip. **(A)** Representative raw and quantified sample images. **(B)** Each box represents one well. Top row: untreated A549 cells. Middle row: non-shRNA control. Bottom row: shPACER knockdown. The blue area highlights the region occupied by cells after the formation of the wound. The orange, red and black areas represent regions of cell growth: 12hr (orange), 24hr (red), 48hr (black). White regions show areas of no growth. The percentage change in growth from 0 to 24 hours was calculated for each sample. **(C)** Analysis of the quantified change in growth area after 24 hours identified significantly less shPACER cell migration (43% Δ) into the wound area compared to shNMC (55% Δ) and wild-type (61%) cells (t(10, n=6) = 2.338, p = 0.0415). Images were captured using the NES Nikon utility at 100 × magnification. Cell migration was quantified using ImageJ and the Wound Healing Size Tool (ImageJ plugin) and adjusted to ensure border areas included all contiguous cells. Error bars represent SEM. Images were cropped for presentation after quantitative analysis. P-values were calculated using a two-tailed unpaired t-test (α = 0.05).

To investigate how PACER affects lung cancer cell invasion, we employed a 3D-Matrigel-based semi-spheroid assay (Figure 4). This approach, adapted from Aslan et al., allows for quantification of invasive outgrowth over several days (Aslan et al. 2021). ShPACER or shNMC cells were grown in standard culture conditions, resuspended and mixed with Matrigel. Matrigel semi-spheres were carefully formed in individual wells (Figure 4A). The semi-spheres grew in complete medium under standard conditions for approximately five days (Figure 4B). Reference images were taken daily to monitor growth and to ensure the semi-spheres remained intact. Growth was measured in pixel area and converted to mm^2^. After day five, the outgrowth areas were measured relative to any slight variations in initial spheroid size (Figure 4C).

**Figure 4.**
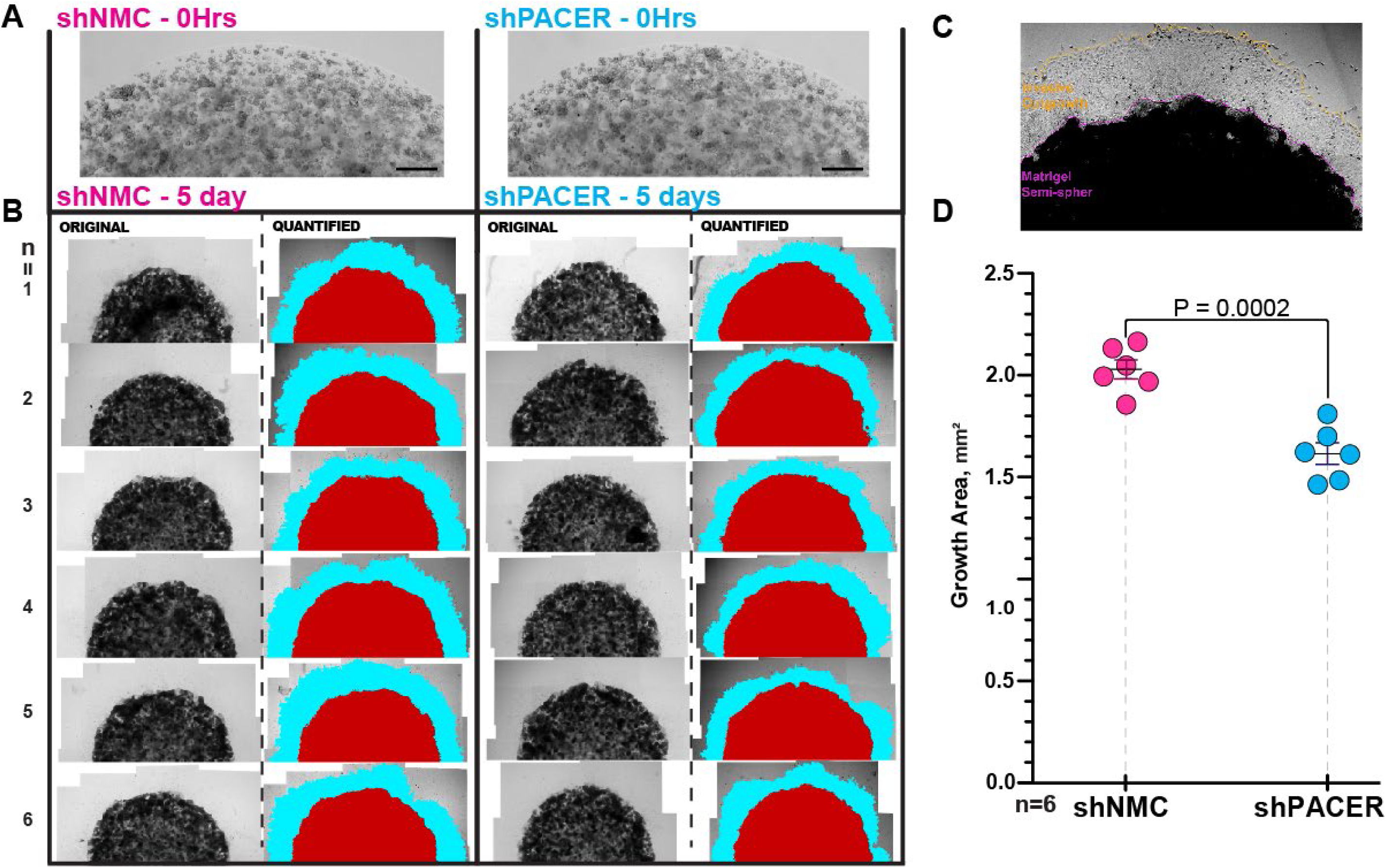
3D-Matrigel Spheroid Invasion Assay. A549 cells transfected with either shNMC (control) or shPACER constructs were embedded in Matrigel semi-spheroids and observed daily for five days, then quantified. Spheroid outgrowth was quantified relative to the initial spheroid sizes to account for slight variations at seeding (<0.5%). **(A)** Original Matrigel spheroids at 0 hrs. The black bar in the bottom right represents a scale of 100 µm. **(B)** and quantified images of spheroids after 5 days. Analyzed using image utilities in ImageJ, the red area corresponds to the spheroid area measured after 5 days, while the cyan areas outline cell outgrowth. **(C)** Reference image showing the image quantification processes. The Matrigel spheroid is outlined in pink while the outgrowth area is outlined in orange **(D).** Quantitative analysis of pixel area showed a decrease (t(10, n=6) = 5.86, p = 0.0002) in the mean total outgrowth area for shPACER spheroids (2.03 mm^2^) vs shNMC (1.61 mm^2^). The central black bar is the mean, and the error bars represent SEM. P-values were calculated using a two-tailed unpaired t-test (α = 0.05).

Quantification of the outgrowth area revealed that shPACER cell growth was reduced compared to shNMC cells, exhibiting a visible difference in growth area (Figure 4D). The initial cell density in the spheroids was consistent, and the area of the initial Matrigel spheroids did not significantly differ between wells. ShNMC cells expanded further from the initial boundaries with an average outgrowth area of 2.03 mm^2^ compared to the shPACER cells with an average area of 1.61 mm^2^. Quantification of spheroid size identified a statistically significant reduction in overall growth area for the shPACER group (t(10, n = 6) = 5.86, p = 0.0002), suggesting a reduction in invasive potential.

Taken together, these findings demonstrate that PACER knockdown significantly impairs both migration and invasion in A549 lung cancer cells. Reduced wound closure and spheroid outgrowth indicate that PACER contributes to a pro-migratory, invasive phenotype, key cellular behaviors associated with metastatic potential in lung adenocarcinoma (Seguin et al. 2022).

### Evolutionary Conservation of the LncRNA PACER in *Mus musculus* and Primates

Given PACER’s influence on inflammatory signaling, cell proliferation, migration and invasion, we next investigated evolutionarily conserved regions and motifs within the PACER transcript that may regulate its expression. PACER, like many lncRNAs, exhibits limited sequence similarity over relatively short spans of evolutionary time (e.g., from the emergence of rodents to the appearance of primates), making it difficult to systematically detect and analyze short, conserved motifs within its sequence.

Here, we utilized LncLOOM, a bioinformatics program developed by the Ulitsky laboratory (LncLOOMv2) (Ross et al. 2021), designed to identify conserved motifs and miRNA binding sites in low-conservation sequences such as lncRNAs. We utilized LncLOOM to analyze the human PACER transcript, six homologous primate transcripts, and the murine homolog Ptgs2os (Table 2). All sequences used in our analysis were downloaded from NCBI. While there is variation between the human PACER transcript and primate sequences, the mouse homolog Ptgs2os is considerably different, consisting of multiple exons spanning approximately 3Kb. The primate sequences were selected based on the quality of their genome annotation and the presence of transcriptional activity near the homologous region.

**Table 2.**
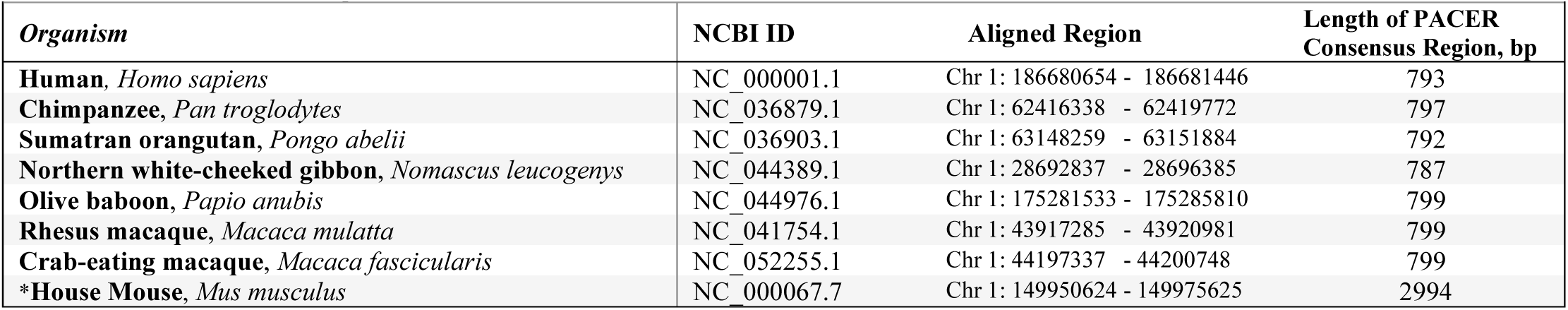
PACER Homologs in Primates & Mice. These organisms in this table were included in the LncLOOM motif conservation analysis of PACER. Excluding *Mus musculus*, a region of approximately 3500bp of the antisense RNA sequence upstream of the 3’ COX-2 start site is downloaded from NCBI for each species and aligned to the human PACER sequence. The aligned PACER consensus region of non-human primates (approximately 800 bp) is used to analyze conserved motifs. \**Mus musculus* is the only species included in our analysis that has an annotated transcript for a PACER homolog, *Ptgs2os*. Ptgs2os has multiple exons, and only the exonic regions were used in our alignment and motif analysis.

When searching between a minimum k-mer length of six and a maximum of fifteen, LncLOOM analysis, paired with *de novo* prediction of miRNA target sites using TargetScan, identified 26 conserved motifs predicted as potential miRNA binding sites. These predicted binding sites were conserved to a depth of seven, meaning they are present in the human and all primate sequences but absent from the mouse homolog (Sup. Table 1). A total of 16 motifs were found to be conserved between the human, primate and mouse homologs (highlighted in gray in Figure 5). Of these 16 motifs, four are predicted to bind miRNAs: miR-18a-5p, miR-196-5p, miR-1306-5p, and miR-423-3p (p = 0.01) (highlighted in color, Figure 5, Sup. Table 1). Interestingly, the predicted miR-423-3p binding site overlaps the distal NF-κB transcription factor binding site present in PACER.

**Figure 5.**
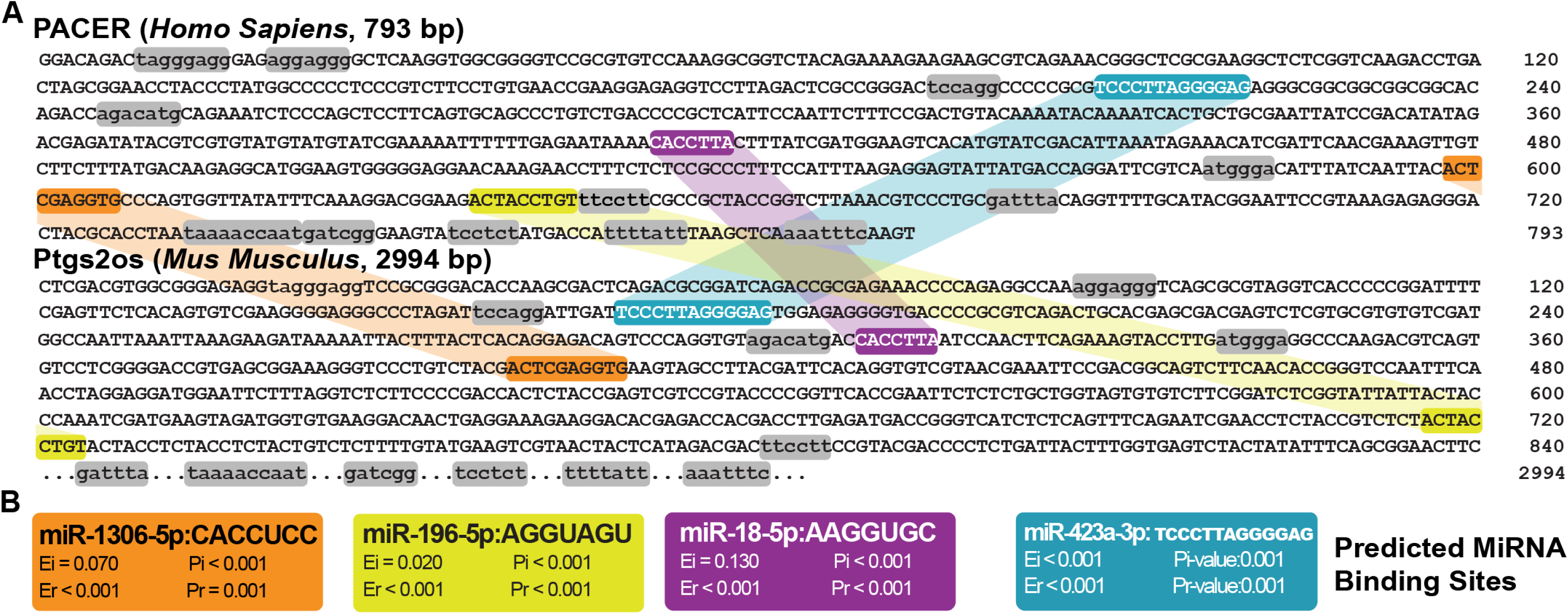
LncLOOM Analysis Identified Conserved MiRNA Binding Motifs Within the LncRNA PACER & Mouse Homolog. LncLOOM is a bioinformatic software developed to identify short, conserved motifs in homologous sequences across diverse species. Human PACER and the homologous regions in seven species were used in this analysis: six primate species and the mouse homolog Ptgs2os (Table 2). **(A)** This figure shows the relationship between the positions of the predicted binding motifs in human PACER and mouse Ptgs2os sequences. A total of sixteen conserved motifs, conserved in all eight homologous sequences, were identified. Motifs highlighted in color are predicted to bind known miRNAs identified by TargetScan (McGeary et al., 2019). **(B)** Predicted miRNA binding sites of the conserved motifs. Motifs are predicted to bind miR-1306-5p, miR-196-5p, miR-18a-5p, and miR-423-3p.

Several conserved motifs are clustered around the 5’ and 3’ domains of the PACER transcript. These clusters may be associated with transcript stabilization as seen in other lncRNAs (Sherpa et al. 2018). There is also a cluster of conserved motifs in the 200-250 bp region flanking a known NF-κB binding sequence and predicted miR-423-3p binding site. These motifs, conserved across evolutionary time, are suggestive evidence of biological importance and indicate that these regions may contain functional domains, such as protein or miRNA binding sites or RNA stability elements.

### In Vivo and Ex Vivo SHAPE-MaP Analysis of PACER in A549 Cells

Establishing that the human PACER transcript harbors multiple motifs and potential regulatory elements conserved across species, we next examined PACER’s overall structure and the structural features of conserved regions in the cellular environment. Chemical probing techniques, such as SHAPE-MaP, are powerful tools for resolving lncRNA secondary structures and linking structural motifs to functional roles. We utilized SHAPE-MaP to probe the structure of PACER in both ex vivo and in vivo conditions. Differences in SHAPE reactivity values, calculated using ΔSHAPE, identified structurally significant motifs that may regulate protein-RNA interactions with PACER. We modeled PACER folding predictions globally using SuperFold, and finally, assessed structural subsets within each condition using the Rsample pipeline to generate and cluster ensemble structures into representative centroid structures.

In vivo and ex vivo SHAPE-MaP analysis was conducted on A549 cells treated with 2A3 or DMSO (vehicle control). Many functional lncRNAs are expressed at low levels (Bubenik et al. 2020). Compared with other RNA species, PACER is less abundant, despite being overexpressed in cancer and inflammatory disease. We addressed complications due to low abundance and the analysis of lncRNAs with complex secondary structures by implementing primer-specific reverse transcription, followed by an overlapping dual-amplicon cDNA amplification strategy (Sup. Figure 2 & Sup. Table 2). This strategy ensured complete coverage of distal regions within 50 bp of the primer-binding sites, overcoming the limitations of the Nextera XT DNA Library Preparation method (Illumina) (Bubenik et al. 2020). Amplicon tiles were designed to be less than 600 bp, within the maximum read length of the NextSeq 2000 (Illumina). Sequencing libraries were prepared from A549 cells treated with either 2A3 or vehicle control (DMSO) and processed separately for in vivo and ex vivo conditions (Figure 6).

**Figure 6.**
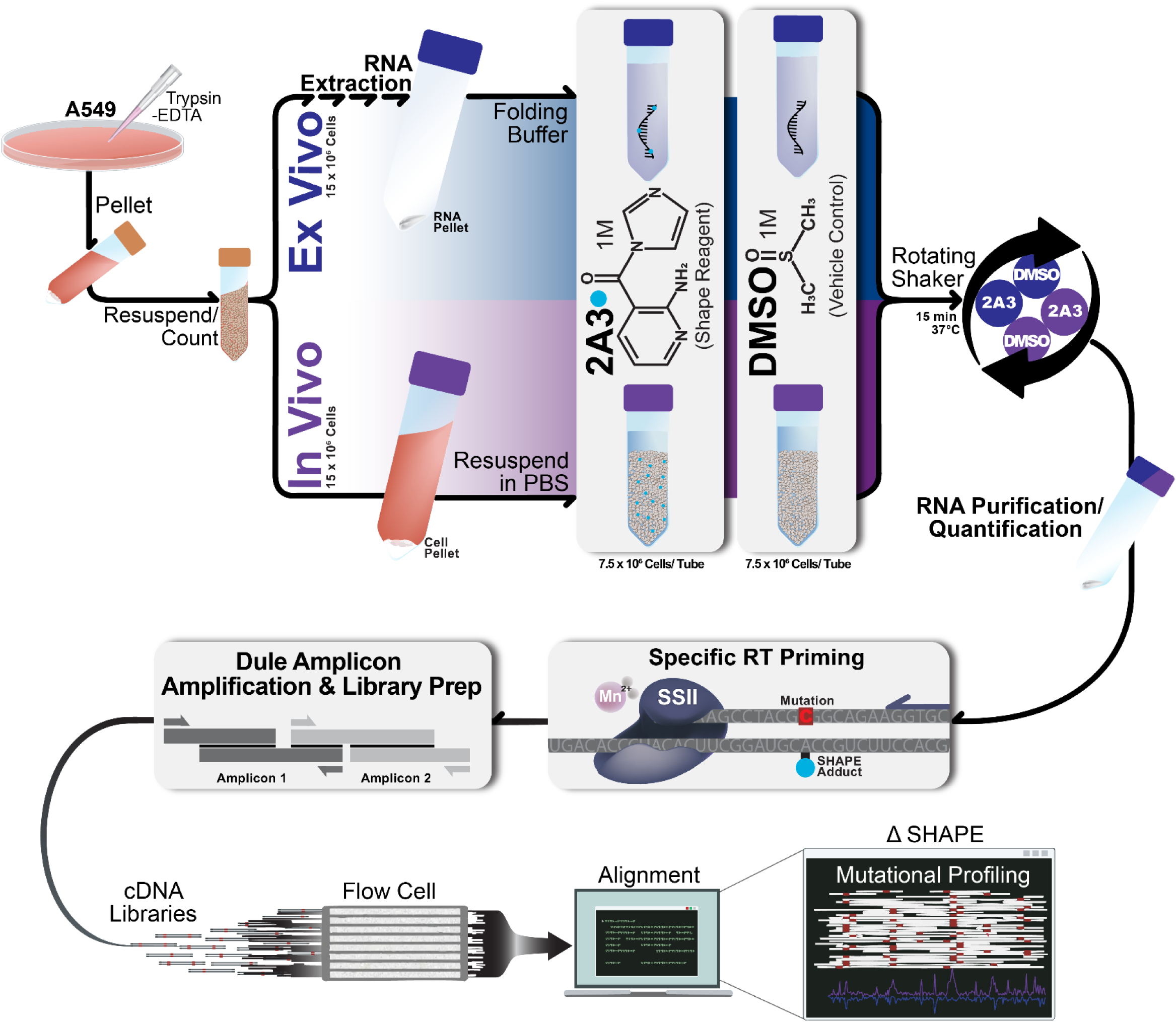
SHAPE MaP Experimental & Analysis Workflow. A549 cells were prepared for in vivo or ex vivo selective 2′-hydroxyl acylation analyzed by primer extension and mutational profiling (SHAPE-MaP). Cells were resuspended, counted and split between ex vivo and in vivo conditions (15 × 10^6^ cells/ condition). For ex vivo probing, RNA was extracted from cultured cells, pelleted, and refolded in folding buffer prior to modification. For in vivo probing, cells were resuspended in phosphate-buffered saline prior to modification. Cells were evenly divided into two treatment groups for ex vivo and in vivo conditions: 2A3 (2-aminopyridine-3-carboxylic acid imidazolide, 1 M) or vehicle control (DMSO, 1 M). Following modification, RNA was isolated, and adducts were encoded during primer-specific reverse transcription using SuperScript II (SSII) in the presence of Mn²⁺. The cDNA was amplified and prepared for sequencing using a dual-amplification, library-preparation strategy. Sequencing reads were aligned to the reference sequence, and mutational profiling was performed to distinguish SHAPE-induced mutations from background (DMSO) error rates. Per-nucleotide mutation frequencies were quantified, background-subtracted, and normalized to produce high-resolution SHAPE reactivity profiles. For each condition, SHAPE reactivity was calculated from n=3 independent biological replicates. These profiles provide a basis for condition-based comparative analyses and secondary-structure modeling.

After initial quality filtering and adapter trimming using the fastp preprocessing tool (Chen 2023), paired-end reads from ex vivo and in vivo libraries were analyzed with ShapeMapper v2.1.5 in amplicon mode. Reads were aligned to the PACER reference sequence (NC_000001.11 Reference GRCh38.p14: 186680654-186681446) using the Bowtie2 Aligner (Langmead & Salzberg 2012) integrated into ShapeMapper. Modified libraries were paired and normalized to vehicle controls for background subtraction. Overall Bowtie alignment rates (Ex vivo: 92.6-96.3%. In vivo: 92.5-96.4%), coverage (100% amplicon coverage for all conditions) and read depth (median per nt read depth: Ex vivo; 1.8 × 10^6^ - 2.2 × 10^6^, In vivo; 1.7 × 10^6^-2.2 × 10^6^) were well above established thresholds, indicating efficient mapping for all libraries (Sup. Figure 3). As expected for highly structured lncRNAs, the fraction of nucleotides with high reactivity was relatively low (Mathews 2004; Smola et al. 2016; Uroda et al. 2019). 3′ and 5′ primer binding sites were masked, and internal primer sites were trimmed and excluded from ShapeMapper analysis. Coverage-weighted global mutation rates confirmed increased mutation frequencies in 2A3-treated samples compared with vehicle controls (Sup. Table 3), consistent with efficient modification.

SHAPE reactivities calculated by ShapeMapper had a higher overall magnitude ex vivo than in vivo. Mean ex vivo reactivity was 0.26 with a 90^th^ percentile of 1.10, whereas the in vivo distribution had a mean of 0.074 and a 90^th^ percentile of 0.58 (Figure 7A). The percentage of nucleotides exceeding established flexibility boundaries was greater in the ex vivo condition (29.4% > 0.3; 21.3% > 0.5; 11.5% > 1.0) than in vivo (15.2%, 8.9%, 5.5%) (Hajdin et al. 2013; Smola et al. 2015b). Correlation of per-nucleotide SHAPE reactivities between ex vivo and in vivo conditions was very low (Pearson’s r = 0.03, Spearman’s ρ = 0.09), indicating that the observed differences are not due to a uniform baseline shift (higher overall magnitude of reactivities in the ex vivo samples).

**Figure 7.**
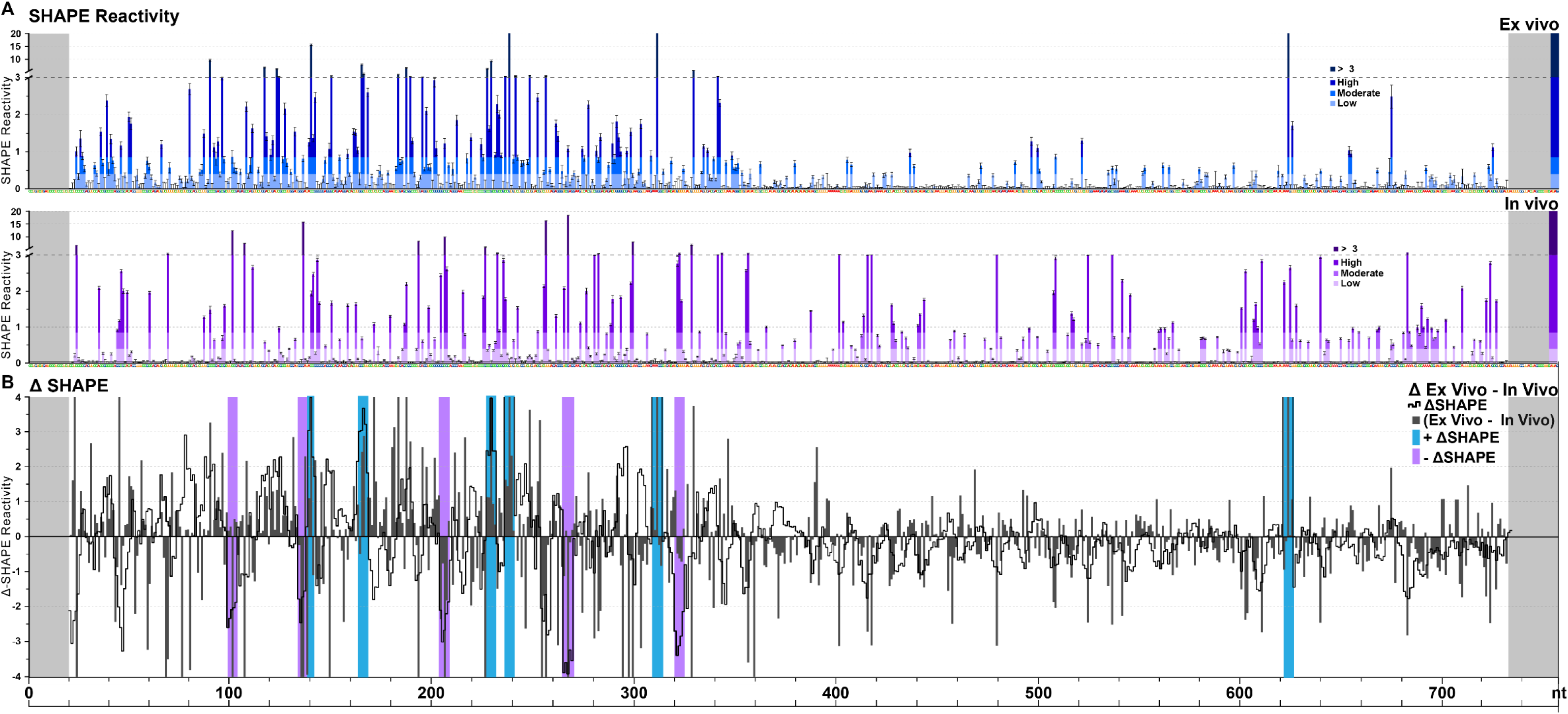
SHAPE-MaP Reactivity Profiles: PACER Ex Vivo & In Vivo Conditions & ΔSHAPE. **(A)** Per-nt SHAPE reactivities (normalized to unmodified) are shown for ex vivo (blue, top panel) and in vivo (purple, middle panel) libraries across the complete PACER amplicon. Gray shaded regions indicate the 5’ and 3’ primer-masked regions excluded from downstream analysis. Internal primer masking excludes internal primer sites from downstream quality control and statistical calculations; however, mutational frequency and reactivity were calculated. Ex vivo PACER exhibited higher mean reactivities (mean 0.26, 90th percentile 1.10) compared to in vivo (mean 0.074, 90th percentile 0.58), with a greater proportion of nucleotides exceeding minimum flexibility thresholds (>0.3: 29.4% vs 15.2%) **(B)** ΔSHAPE (ex vivo - in vivo) SHAPE reactivity values (black line), raw ex vivo - in vivo values (grey bars). Blue and purple shaded bars represent sites protected in vivo (blue, + ΔSHAPE) or more reactive in vivo (purple, - ΔSHAPE) compared to ex vivo. A region was considered significant where for ≥ 3 of 5 consecutive nucleotides, non-overlapping 95% confidence intervals between modified and control libraries as determined by the Z-factor test (Z > 0), and a standard score threshold of |S| ≥ 1.

### Modular Domain Organization of PACER Revealed ΔSHAPE and SuperFold Analysis

We measured variation between ex vivo and in vivo reactivity across the PACER transcript using the ΔSHAPE method, as described by Smola et al. (Smola et al. 2015a). ΔSHAPE was defined as ex vivo minus in vivo SHAPE reactivity, such that positive values identify areas of in-cell protection and negative values indicate increased reactivity in cells (Figure 7B). Statistical thresholds were set as described by Smola et al. with per-nucleotide differences filtered by a Z-factor test enforcing non-overlapping 95% confidence intervals (Z > 0), and standard score criteria (|S| ≥ 1) (Smola et al. 2015a). Regions were considered significant where ≥ 3 of 5 consecutive nucleotides met both criteria.

ΔSHAPE analysis identified 11 significant regions of differential reactivity between ex vivo and in vivo conditions, comprising six sites that were more protected in vivo (+ΔSHAPE) and five sites that were more reactive in vivo (-ΔSHAPE). Protected and reactive sites were present in multiple locations across the transcript, but several clustered between nts ∼200-400. Additional sites appeared towards the 5′ (nt 100) and 3′ (nt 624) ends. All sites are summarized by the blue (protection sites, +ΔSHAPE) and purple (enhancement sites, -ΔSHAPE) shaded regions in Figure 7B. ΔSHAPE results indicate that the in vivo transcript has several discrete protected regions and areas that undergo dynamic structural shifts, consistent with external binding interaction and formation of ribonucleoprotein (RNP) complexes.

We utilized SuperFold, which combines SHAPE data with thermodynamic folding to infer consensus base-pairing models and Shannon entropy profiles. This analysis enabled direct comparison of RNA structural models across ex vivo and in vivo conditions. Overall base-pairing fractions were comparable across conditions (64.0% ex vivo vs. 63.2% in vivo), but there were clear differences in the pairing probabilities (Ppair) and length of interactions (Figure 8A). Ex vivo PACER contained more total arcs above the probability threshold (Ppair ≥ 0.10, 269 ex vivo vs. 120 in vivo) but maintained a similar mean pairing probability (0.51 vs. 0.50 in vivo) and percentage of short to midrange helices (≤ 200 nt, 86.6 for both in vivo and ex vivo). The in vivo and ex vivo transcripts had the same percentage of long-range bps (≥ 200, 13.3%), and the in vivo transcript had a marginally higher level of very long-range interactions (≥ 400, 5.2% vs. 6.6% in vivo). While the percentage of very long-range interactions was similar, in vivo mean and 99^th^ percentile paring probabilities were substantially higher (mean Ppair = 28.4% vs. 50.1% in vivo, 99^th^ percentile = 0.360 vs. 0.866 in vivo). These data indicate that, despite similar overall pairing, the in vivo transcript resolves into fewer, higher-probability long-range contacts indicative of a more stabilized architecture.

**Figure 8.**
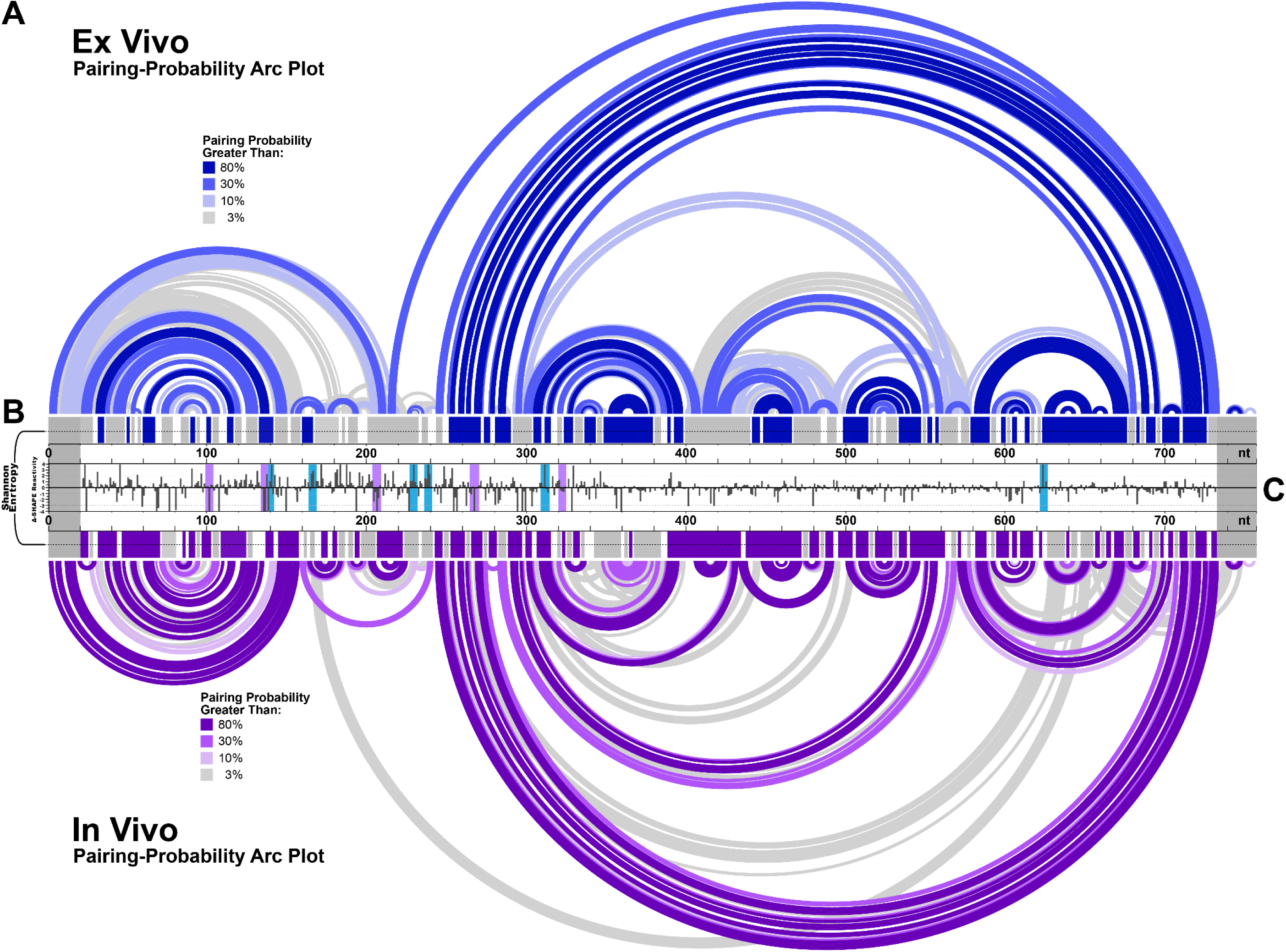
SuperFold Analysis of PACER Secondary Structure Paired With ΔSHAPE. **(A)** SuperFold base-pairing arc plots comparing ex vivo (top) and in vivo (bottom) conditions. Overall base-pairing fractions were similar (ex vivo 64.0%; in vivo 63.2%). Ex vivo PACER had more arcs with Ppair ≥ 0.10 (269 ex vivo vs. 120 in vivo), with a similar mean pairing probability (0.51 vs. 0.50 in vivo) and the same proportion of short-to-midrange helices (≤200 nt: 86.6% in both). The proportion of long-range base pairs (≥200 nt) was the same in both conditions (13.3%), whereas very long-range pairs (≥400 nt) were slightly more frequent and of higher confidence in vivo (6.6% vs. 5.2% ex vivo; 99th percentile Ppair 0.866 vs. 0.360). **(B)** Shannon entropy profiles for ex vivo (blue) and in vivo (purple). In vivo PACER exhibited reduced entropy (mean 0.131 vs. 0.196 ex vivo) and extended low-entropy tracts, consistent with a more compact and stable domain organization. Grey shaded regions indicate high-entropy intervals (>0.15 for ≥5 consecutive nts), while colored bars mark low-entropy intervals (≤0.15 for ≥5 consecutive nts). **(C)** ΔSHAPE (ex vivo - in vivo) SHAPE reactivity overlay showing significant differences between conditions. Blue shaded bars mark regions protected in vivo (+ΔSHAPE), purple shaded bars mark regions more reactive in vivo (−ΔSHAPE). Six ΔSHAPE windows overlapped with low-entropy tracts, identifying sites of structural remodeling in the cellular context. Together, these data support a model in which PACER adopts a stabilized 5′ domain and more flexible central and 3′ domains, consistent with modular organization and potential RNP stabilization in vivo.

Shannon entropy profiles provided complementary evidence of condition-specific folding stability. In vivo entropy values were noticeably reduced (mean 0.131, median 0.110) relative to ex vivo (mean 0.196, median 0.148), with a greater fraction of nucleotides classified into stable low-uncertainty bins (< 0.15: 50.5% vs. 64.9% in vivo). Shannon profiles result in extended tracts of low-entropy structure in vivo, in contrast to the fragmented organization observed ex vivo. The Shannon entropy profiles indicate that the in vivo transcript favors a more consistent and predictable architecture than the ex vivo condition.

Overlaying ΔSHAPE and Shannon entropy profiles further highlighted condition-dependent remodeling often linked to function. We investigated regions with low ΔSHAPE reactivity and low Shannon entropy, which are known to be correlated with high structural consistency and functionality (Smola et al. 2015b). We identified overlaps between ΔSHAPE sites and low Shannon entropy regions (≥ 3 consecutive nts with entropy ≤ 0.15) as shown in Figure 8B-C. Six of eleven ΔSHAPE windows overlapped with low-entropy tracts. Sites with higher reactivity in vivo (-ΔSHAPE) were located in ex vivo low-entropy helices, consistent with helix destabilization in cells. In contrast, sites with reduced reactivity in vivo (+ΔSHAPE) overlapped low-entropy helices in vivo, consistent with helix stabilization or potential RNP protection (Figure 9). Several sites (nts 100-102, 135-137, and 266-268) that formed well-defined helices ex vivo became more reactive in vivo, indicating that these regions undergo helix destabilization or loop opening inside cells. Conversely, regions with reduced reactivity in vivo (+ΔSHAPE) overlapped with low-entropy structural motifs, suggesting helix stabilization and possible involvement of RNP interactions. The clustering of ΔSHAPE transitions with distinct low-entropy tracts, along with the enrichment of stable helices at the 5′ end relative to the more variable central and distal regions, supports a modular organization of PACER, in which separate domains have discrete roles in RNA interaction and structural maintenance.

**Figure 9.**
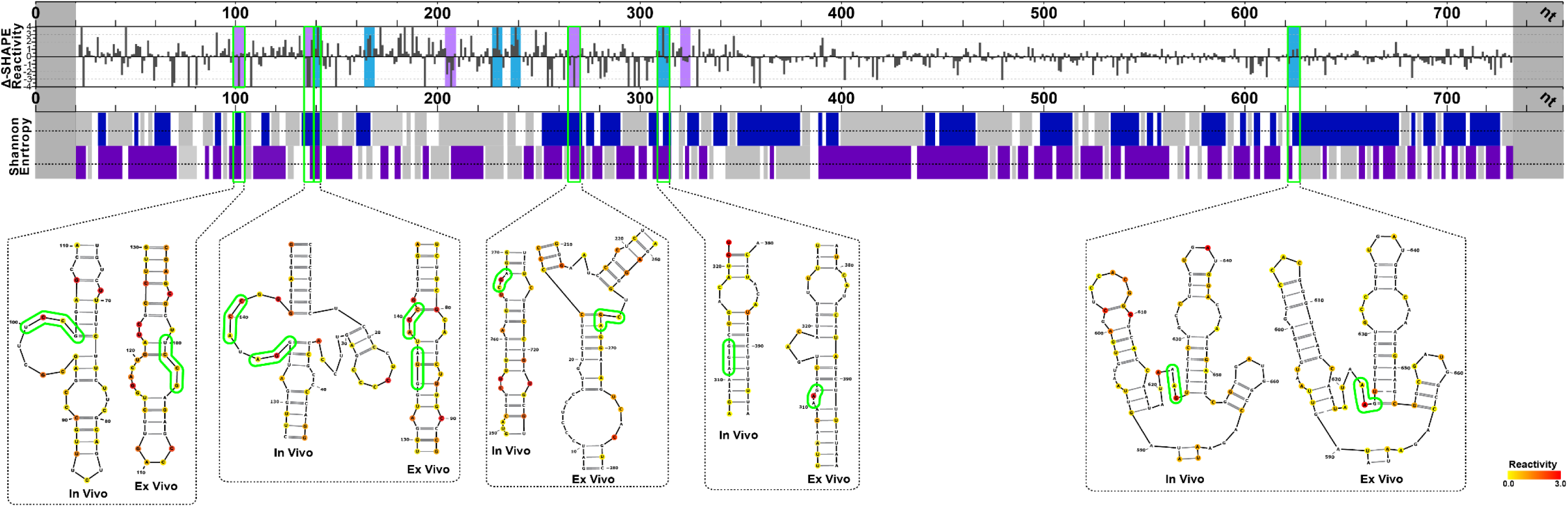
Superfold Local Structural Comparison. Local structural elements for regions of the PACER lncRNA with overlapping significant ΔSHAPE and low Shannon entropy windows. The ΔSHAPE (ex vivo - in vivo) SHAPE reactivity plot (outlined in Figure 7) was overlaid with Shannon entropy plots (outlined in Figure 8) to identify regions that are more protected or more reactive in vivo. Comparison of ΔSHAPE and Shannon entropy revealed that six of the eleven ΔSHAPE windows overlapped with low-entropy tracts. Sites with increased reactivity in vivo (−ΔSHAPE) coincided with helices that were structured ex vivo, whereas sites with reduced reactivity in vivo (+ΔSHAPE) overlapped stable low-entropy helices in vivo (outlined in green).

### Ensemble Structure Clustering and Comparison of In Vivo and Ex Vivo PACER

To characterize the ensemble diversity of PACER secondary structure within each condition, we utilized the Rsample stochastic sampling and clustering pipeline (Reuter & Mathews 2010). This approach generates a large set of SHAPE-constrained ensemble structures (n=1000, per condition), applies hierarchical clustering, and defines representative centroid structures that illustrate the conformational heterogeneity not detectable by global folding algorithms such as SuperFold.

Rsample clustering revealed that the in vivo ensemble was dominated by two major centroid structures, IV-C1 (38.2%) and IV-C2 (58.9%), with a negligible IV-C3 cluster (2.9%) (Figure 10A). By contrast, the ex vivo ensemble was heavily weighted towards a single cluster, with EX-C1 (73.6%) as the primary conformation and EX-C2 (24.4%) as a minor secondary centroid, with the remaining clusters accounting for a negligible 2.0% of structures. Each centroid structure contained approximately 200 bps comprising 40-50 structured helices. Base-pair consensus analysis showed that both IV-C1 and IV-C2 aligned more closely with EX-C2 (Figure 10A-B, sharing 136 bps; Jaccard (*J*) = 0.562 for IV-C2/EX-C2) than with EX-C1 (*J* = 0.27-0.32 for IV-C1/C2). Thus, there is a condition-dependent shift in structural equilibrium with in vivo PACER favoring IV-C1 and IV-C2 (EX-C2 like) conformations, while ex vivo is dominated by EX-C1.

**Figure 10.**
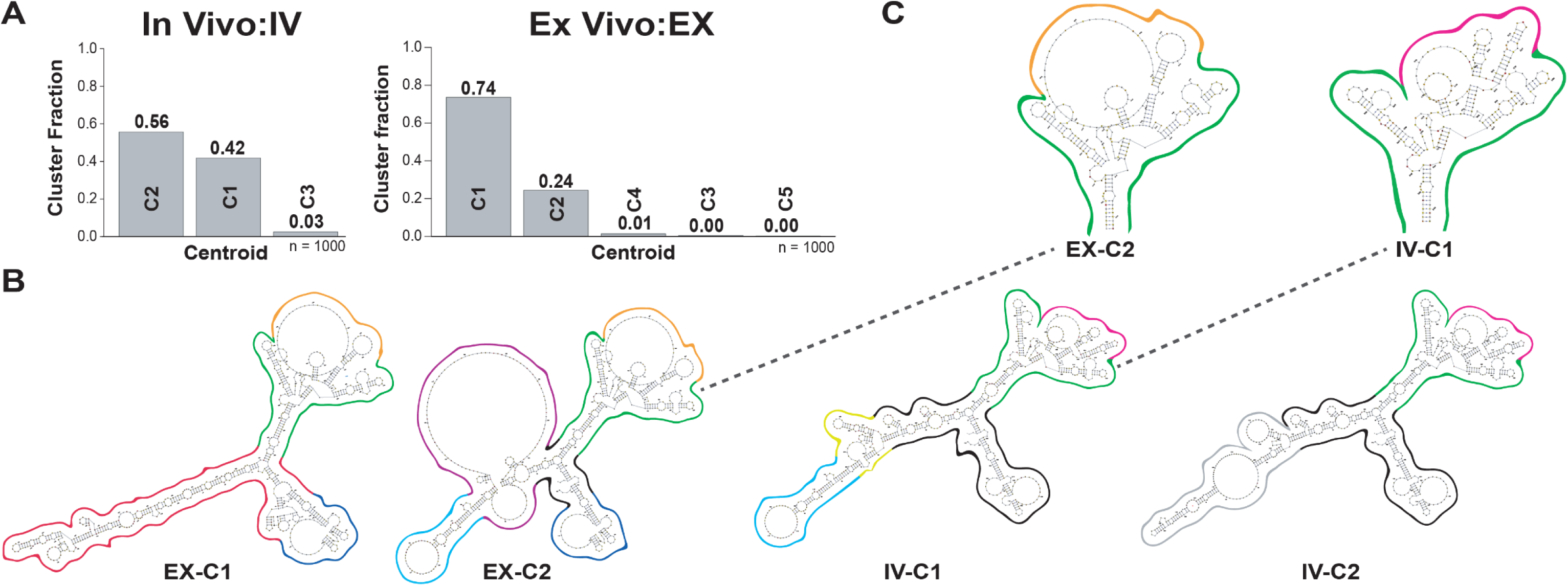
Rsample Ensemble Clustering of PACER Secondary Structures. **(A)** Hierarchical clustering of SHAPE-constrained structures (n = 1,000 per condition. The in vivo ensemble was partitioned into two dominant centroids, IV-C1 (38.2%) and IV-C2 (58.9%), with a minor IV-C3 (2.9%). The ex vivo ensemble was weighted toward a single dominant centroid, EX-C1 (73.6%), with EX-C2 (24.4%) as a secondary structure. **(B)** Base-pair consensus comparison of centroids. Conserved base pairs shared across centroids (black outline) highlight a stable structural core. Both in vivo centroids aligned more closely with EX-C2 (Jaccard J = 0.562 for IV-C2/EX-C2) than with EX-C1 (J = 0.27-0.32), indicating structural reorganization between conditions. Two major domains were identified: nts 50-300 (variable across centroids) and nts 310-635 (broadly conserved but containing a flexible tract at nts 455-550). Colored overlays denote condition-specific pairings (e.g., IV-C1, yellow; IV-C2, grey; EX-C1, red; EX-C2, purple). **(C)** Structural variability in the 310-635 nt domain, consistent across centroids but containing many low probability pairs, suggests regional flexibility despite overall conservation. Together, these results support a modular architecture with a stabilized 5′ domain and more dynamic central and distal domains.

Rsample ensemble clustering showed that the conformation of PACER is condition-dependent, with two dominant in vivo centroids and a single, more predominant ex vivo state. Direct comparison of pairing and structural motif conservation identified a set of conserved base pairs (Figure 10B, black outline) present in all centroids except EX-C1, as well as two domains of interest spanning nts 50-300 and nts 310-635 (green outlines). Within the 5′ domain, the 50-240 nt region exhibited unique pairing patterns between in vivo centroids, with EX-C2 showing consistent overlap with IV-C1 from nts 70-160 (IV-C1/EX-C1, light-blue; IV-C2, grey). The adjacent segment (160-300 nt) showed no consistent fold across centroids, with each adopting distinct helix arrangements (Figure 10B, IV-C1, yellow; IV-C2, grey; EX-C1, red; EX-C2, purple). The larger 310-635 nt domain displayed broad structural conservation across centroids but contained a nested tract (455-550 nt) that was internally consistent within conditions (orange for EX-C1/C2; pink for IV-C1/C2) yet differed between ex vivo and in vivo ensembles. While this region (nts 310-635) was conserved across all centroids, it contains many unbound, structurally flexible motifs, consistent with the SuperFold analysis (Figure 10C). Thus, PACER features a conserved central domain (nts 310-635) with condition-specific remodeling and a variably folded 5′ module (nts 50-300), making EX-C1 the principal outlier.

These findings indicate that PACER adopts a more compact in vivo conformation, with a modular architecture characterized by a stabilized 5′ domain and a more dynamic central and 3′ domain. While certain regions become destabilized in vivo, increasing local flexibility, others remain protected within stable helices, a pattern consistent with potential stabilizing interactions (Figure 11A-B). Although additional studies are needed to directly link these structural features to PACER’s regulatory function, the overlap between variable regions and low-entropy domains highlights candidate functional sites and supports further mechanistic investigation.

**Figure 11.**
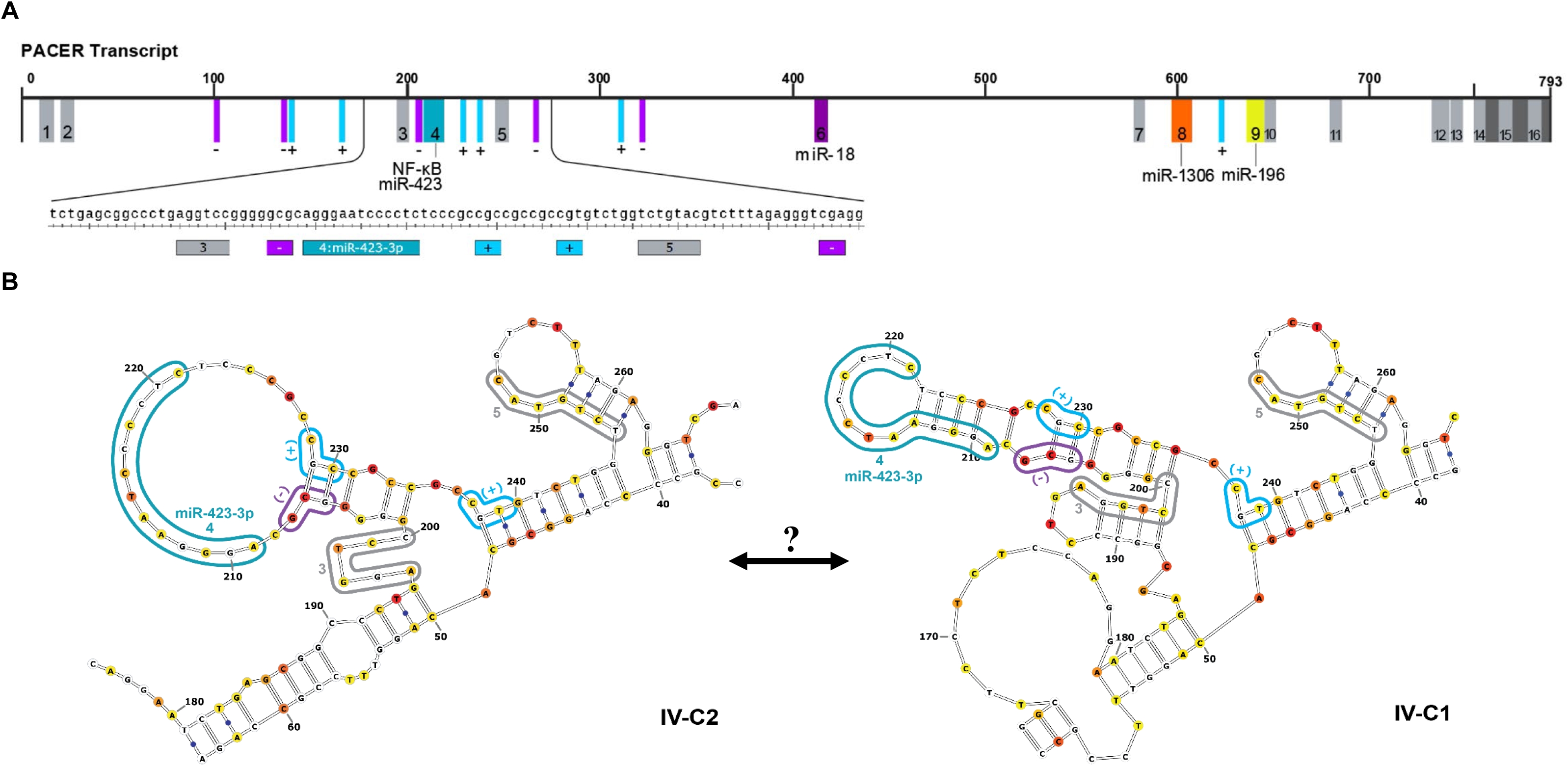
Conserved Motifs and Significant ΔSHAPE Regions in PACER. **(A)** Conserved sequence motifs (gray boxes, numbered 1-16) identified by LncLOOM analysis. Colored boxes indicate predicted miRNA binding motifs identified by TargetScan: teal, miR-423-3p/ NF-κB binding site (nts 209-221); dark purple, miR-18a-5p (nts 412-418); orange, miR-1306-5p (nts 598-607); and yellow, miR-196-5p (nts 637-651). Overlaid ΔSHAPE regions denote nucleotides exhibiting differential reactivity between ex vivo and in vivo conditions: light blue (+ΔSHAPE) marks sites protected in vivo; magenta (-ΔSHAPE) marks regions more reactive in vivo, indicating localized helix destabilization or enhanced flexibility. Below the main track is a sequence level view of the 181-270 region containing the known NF-κB binding site, predicted miR-423-3p motif. **(B)** LncLOOM motifs and ΔSHAPE regions mapped to the IV-C2 and IV-C1 structural determinations derived from Rsample clustering. This region illustrates the alternative structural arrangements between IV-C1 and IV-C2 surrounding the NF-κB/ miR-423-3p predicted interaction domains, suggesting condition-dependent conformations.

## Discussion

LncRNAs play diverse roles in cancer biology, but their mechanisms often remain incompletely defined. In this study, we examined the lncRNA PACER, integrating phenotypic assays, evolutionary analysis, and SHAPE-MaP RNA structural probing. These combined approaches allowed us to evaluate both the functional impact of PACER on COX-2 signaling and the structural features underlying its ability to regulate COX-2 mRNA expression. Our combined analysis supports a model in which PACER promotes cytokine-driven COX-2 transcription, reinforcing its role in promoting LUAD cell proliferation, migration, and invasion. Additionally, we identified structural features that may give rise to PACER’s ability to regulate COX-2 mRNA expression.

Stable shRNA knockdown experiments reduced PACER expression by greater than 50% and decreased COX-2 mRNA by nearly two-thirds in A549 cells (Figure 2B-C). Upon TNF-α/IL-1β stimulation, WT and shNMC cells upregulated PACER and COX-2, whereas shPACER cells showed a blunted response, indicating that PACER is necessary to achieve robust NF-κB-mediated COX-2 activation (Figure 2D). These findings suggest that PACER has a role in amplifying cytokine-induced NF-κB signaling and sustains the COX-2/PGE₂ feedback loop that drives chronic inflammatory and proliferative pathways (Castellone et al. 2005; Filippelli et al. 2024). Consistent with a canonical decoy model (Zhang et al. 2020), PACER sequesters inhibitory NF-κB p50 to enable promoter activation, while PGE₂ stimulation further enhances PACER expression, reinforcing the self-perpetuating PACER/COX-2/PGE₂ regulatory circuit (Desind et al. 2022; Krawczyk & Emerson 2014).

The proliferation assays show that PACER knockdown reduced A549 cell growth by ∼24% at Day 4 (Figure 2E). This decrease in proliferation likely reflects attenuation of COX-2-driven autocrine pathways, possibly mediated through PGE₂-dependent EP2 and EP4 receptors, which activate downstream β-catenin/PKA and ERK/AKT cascades to sustain proliferative signaling (Filippelli et al. 2024; Finetti et al. 2020; Jara-Gutierrez & Baladron 2021). Notably, our results show that even an incomplete knockdown of PACER expression is sufficient to disrupt proliferative signaling, suggesting PACER’s importance in maintaining COX-2-driven tumor growth in LUAD.

Beyond proliferation, COX-2 metabolites, mainly PGE₂, are also well-established drivers of cancer cell motility and invasion, acting through EP4 receptor-mediated β-arrestin1/c-Src signaling and EGFR-dependent activation of Akt (Buchanan et al. 2003; J. I. Kim et al. 2010). Wound-healing assays demonstrated that PACER knockdown was sufficient to significantly impair the migratory capability of A549 cells, consistent with the idea of PACER as an important regulatory component of COX-2 signaling (Figure 3A-C). Under low-serum (1%) conditions designed to minimize proliferation effects, shPACER cells covered only 43% of the wound area at 24 hours compared with 55% coverage for shNMC and 61% for WT controls (Figure 3C). Our findings indicate that PACER contributes to the pro-migratory phenotype of LUAD cells, extending its influence beyond proliferative growth to additional hallmarks of metastasis.

To complement our two-dimensional migration assays, we assessed LUAD cell invasion using a 3D Matrigel semi-spheroid assay, which recapitulates the physical architecture of the extracellular matrix and cell-cell interactions characteristic of solid tumors (Aslan et al. 2021; Vinci et al. 2015) (Figure 4). Importantly, the matrix resistance and pressure required for cells to organize into multicellular tumoroids are not captured by 2D wound-healing assays, highlighting the physiological relevance of 3D models for assessing metastatic potential. PACER knockdown significantly reduced invasive outgrowth compared with shNMC controls, consistent with our proliferation and wound-healing assays. The reduced outgrowth of shPACER cells from the Matrigel spheroids was visibly apparent, indicating that PACER knockdown reduced not only proliferative and migratory behaviors but also the ability of LUAD cells to invade through extracellular matrix-like environments. Together, we show that knockdown of PACER expression attenuates basal and cytokine-induced COX-2 signaling, and our phenotypic assays demonstrate that reduced PACER expression leads to reduced proliferation, migration, and invasion in LUAD cells. Given PACER’s established regulation of COX-2 through p50 sequestration, this research provides strong support for the relevance of PACER as both a biomarker and therapeutic target.

Together, our findings demonstrate that PACER knockdown significantly attenuates both basal and cytokine-induced COX-2 signaling, leading to clear reductions in LUAD cell proliferation, migration, and invasion. Combining these phenotypic outcomes with established roles in regulating COX-2 transcription through NF-κB p50 sequestration underscores PACER’s potential as both a mechanistic biomarker of COX-2 pathway activation and a therapeutic target in inflammatory cancers.

To explore how PACER functions, we examined PACER conserved regulatory elements using LncLOOM, a comparative tool for detecting and analyzing motifs in low-homology transcripts (Ross et al. 2021). Our LncLOOM analysis identified multiple short, deeply conserved motifs in PACER shared across primates and the murine homolog Ptgs2os (Sup. Table 1), indicating functional significance despite limited overall sequence conservation. Four motifs overlapped predicted miRNA binding sites (miR-18a-5p, miR-196-5p, miR-1306-5p, and miR-423-3p; Figure 5), suggesting that PACER may be regulated by miRNAs or act as a competing endogenous RNA. Notably, the predicted miR-423-3p binding site overlapped with PACER’s NF-κB domain, introducing the potential for competitive interactions. Such decoy-style interactions are common in lncRNA-miRNA networks. MiR-423-3p is overexpressed in lung adenocarcinoma and other inflammatory cancers. It promotes proliferation, invasion, and EMT by repressing tumor suppressors, including CYBRD1 and p21. (Li et al. 2015; Ma et al. 2021; R. Wang et al. 2019).

Conserved motifs cluster at PACER’s 5′ and 3′ domains and near the 200-250 nt region flanking the NF-κB/miR-423-3p site (Figure 11). The possibility of links to other components of transcriptional machinery provides complexity to the PACER’s regulation and the already complex transcriptional landscape of COX-2.

Using SHAPE-MaP and SuperFold, we generated the first secondary structure of PACER and characterized changes in architecture between ex vivo and in vivo conditions to identify regions likely involved in higher-order interactions and structural stability (Figure 7-11). SHAPE-MaP analysis revealed that PACER adopts a more compact, stabilized architecture in vivo, with a discrete, ordered 5′ domain. Lower SHAPE reactivities (mean in vivo: 0.074 vs. ex vivo 0.260) and reduced Shannon entropy (mean in vivo: 0.131 vs. ex vivo 0.196). Thus, PACER is structurally more constrained in vivo, having extended low-entropy tracts consistent with the formation of these stabilized domains.

SuperFold modeling indicated that PACER exhibits a stable 5′ domain and a flexible central and 3′ domain. SuperFold analysis also showed that PACER contained fewer arcs in vivo (269 ex vivo vs. 120 in vivo, Ppair ≥ 0.10) but had higher probability arcs longer than 400 nt (mean Ppair = 28.4% vs. 50.1% in vivo, 99^th^ percentile = 0.360 vs. 0.866 in vivo). Many of the high-probability long-range pairs observed in vivo fall in low-entropy regions (Figure 8) and flank conserved regions identified during our LncLOOM analysis. Our ΔSHAPE results were consistent with our Superfold analysis, supporting stability in the 5′ domain and a more flexible central region and patterns observed in large lncRNAs stabilized by ribonucleoprotein interactions (Frank et al. 2020; Lin et al. 2018; Sherpa et al. 2018; Somarowthu et al. 2015). This evidence supports a model in which PACER’s in vivo architecture is organized into discrete functional modules. Modular architectures have been described for several long lncRNAs, including HOTAIR (∼2.2 kb), MEG3 (∼1.6 kb), and other lncRNAs of comparable length, such as the 632 nt isoform of GAS5 implicated in multiple cancers (Frank et al. 2020; Sherpa et al. 2018; Somarowthu et al. 2015; Xu et al. 2019; Zhou & Chen 2020). NEAT1 also exhibits similarities, where long-range 5′-3′ base pairing flank regions known to be associated with protein interactions important to its function (Lin et al. 2018). Long-range stabilization of the 5′ domain of PACER may similarly define functional regions necessary for interactions with NF-κB p50 and transcriptional regulation of COX-2.

Rsample ensemble analysis revealed a modular architecture with a conserved structural core and a region spanning the central and 3′ domains, which was preserved across all predicted centroids. In vivo, PACER formed two dominant centroids (IV-C1, 38.2%; IV-C2, 58.9%) compared to a single structure ex vivo (EX-C1, 73.6%), indicating condition-dependent flexibility rather than transcript-wide reorganization. While SuperFold analysis revealed limited structural consensus within the central domain (nts 400-510), Rsample showed a broader conserved region (nts 310-635) containing a variable subregion (nts 455-550) that was structurally consistent within each condition but differed between them (Figure 10C). This subregion appeared flexible ex vivo yet displayed low SHAPE reactivity, consistent with transient or heterogeneous base pairing that obscures modification without forming a single stable structure. In vivo, the same region adopted a more stable architecture, indicative of stabilizing protein- or RNA-mediated interactions. Notably, conserved motifs including miR-18a-5p and miR-1306-5p binding motifs were located in the stable flanking regions, but not within the variable region.

Although inherent differences between SuperFold and Rsample models were anticipated, we observed condition-dependent structural changes localized to the 5′ domain (nts 50-300). This is consistent with our ΔSHAPE and SuperFold data and coincides with key interacting motifs, including the NF-κB domain/ predicted miR-423-3p binding site (Figure 11A). Conserved motifs identified by LncLOOM and ΔSHAPE-defined regions flanked the predicted NF-κB/miR-423-3p interaction domain, which resides within a hairpin spanning nts 206-229 in IV-C2 (Figure 11B). This hairpin is distinctly rearranged relative to IV-C1. Collectively, the presence of multiple known and punitive interaction sites and evidence of structural rearrangement indicate that conformational switching may serve as a regulatory mechanism controlling protein and miRNA accessibility within this region.

This study presents the first structural and functional characterization of the lncRNA PACER in LUAD. Our findings support a model in which PACER enhances basal and cytokine-induced COX-2 transcription, reinforcing downstream pathways that promote proliferation, migration, and invasion. SHAPE-MaP and Rsample analyses revealed a modular architecture with a conserved, structured 5′ domain stabilized by long-range interactions and a more flexible central and 3′ domain. These structural modules coincide with conserved motifs identified by LncLOOM and ΔSHAPE, including regions associated with NF-κB p50 and predicted miRNA interactions. Together, the data suggest that dynamic conformational changes of key motifs may modulate PACER’s interactions with proteins and miRNAs. This work establishes PACER as a key regulator of COX-2 signaling in inflammatory cancers, such as LUAD and provides a structural framework for its exploration as a biomarker and therapeutic target in inflammatory cancers.

## Materials and Methods

### Cell Culture

A549 cells (ATCC, Manassas, VA) were grown in complete medium, Dulbecco’s Modified Eagle Medium (DMEM, MilliporeSigma, St. Louis, MO) supplemented with 10% fetal bovine serum (FBS, R&D Systems, Minneapolis, MN), 2% glutamine (Corning, Corning, NY) and 1% penicillin/streptomycin (Corning). Cells were incubated at 37 °C in 5% CO_2_ incubator (standard conditions) for all cell culture experiments unless otherwise indicated. For passaging and subculturing for experiments, cells were grown in a 100mm culture dish to approximately 75-80% confluency, washed with phosphate-buffered saline (PBS) (Thermo Fisher, Gibco, Waltham, MA) and resuspended by adding 1 mL of 0.05% trypsin-EDTA (Corning) and incubated in standard conditions for 6 minutes before replating in fresh medium. The TC10 Automated Cell Counter (Biorad, Hercules, CA) was used to count and quantify cells. Only cells with fewer than 18 total passages were used for experiments.

### RNA Extraction

RNA was extracted using TRIzol (Thermo Fisher Scientific) and phenol-chloroform phase separation. 200 µL of phenol-chloroform was added to the cells after they were lysed in TRIzol, then shaken for 15 seconds, incubated for 3 minutes at room temperature, and centrifuged (12,000 × g, 15 minutes, 4 °C). The aqueous phase was transferred to new 1.5 mL tubes, mixed with 500 µL 100% isopropanol, incubated 10 min at room temperature, and centrifuged (12,000 × g, 10 min, 4 °C). Supernatant was discarded, and pellets were washed twice with 1 mL 75% ethanol, centrifuged (7,600 × g, 4 °C), and dried. RNA pellets were resuspended in molecular-grade water and quantified using the NanoDrop One Spectrophotometer (Invitrogen).

### Quantitative Real time-PCR

For both PACER and COX-2 RNA analysis, complementary DNA (cDNA) was synthesized using the Qiagen QuantiTect Reverse Transcription kit (Qiagen) following the standard protocol. cDNA preparations contained 1µg of total RNA. qRT-PCR was performed using the CFX96 Real-Time PCR system (Bio-Rad). GAPDH served as a normalizing control. The following primers (synthesized by: Millipore Sigma, PACER; Integrated DNA Technologies, GAPDH; Qiagen, COX-2) were used to detect and quantify RNA expression:

PACER forward 5′-TGTAAATAGTTAATGTGAGCTCCACG-3′
PACER reverse 5′-GCAAATTCTGGCCATCGC-3′
GAPDH forward 5′-CCACCCATGGCAAATTCCATGGCA -3′
GAPDH reverse 5′- TCTAGACGGCAGGTCAGGTCCACC-3′
COX-2 QuantiTect Primer Assay (NM_000963, Qiagen)

Amplification was performed using QuantiTect SYBR Green Master Mix (Qiagen). No template controls and cDNA preps containing no reverse transcriptase were included to ensure samples were not contaminated. The following cycling conditions were used: (1) 94°C for 3 min, (2) 40 cycles of 94°C for 15 sec, 55°C for 30 sec, 68°C for 30 sec (collection step). Initial validation, including melt curve analysis and electrophoresis of amplified products, was performed. -Log_2_ comparative CT (2^-ΔΔCT^) analysis was used to quantify gene expression changes relative to the normalization control GAPDH.

### Lentiviral Knockdown of PACER

A549 cells were transduced with lentiviral particles carrying shRNAs targeting PACER lncRNA to generate a stable knockdown model (Sup. Figure 1). Two candidate shRNAs (Table 1) were cloned into the pLKO.1-puro vector (shPACER, MilliporeSigma); a non-targeting shRNA vector (pLKO.1-puro-Non Mammalian, MilliporeSigma) served as control (shNMC). Lentivirus was produced by co-transfecting HEK293T cells (ATCC) with target or control vectors and packaging/envelope plasmids (gift of Dr. Utz Herbig, Rutgers NJMS) using Lipofectamine 3000 (Thermo Fisher). Medium was collected at 48 and 72 hours, pooled, and concentrated by centrifugation.

A549 cells (3.0 × 10^4^/ well, 96-well plate) were seeded in antibiotic-free DMEM with 10% HI-FBS and 2% glutamine and incubated overnight (∼70% confluency). Polybrene (8 µg/mL, MilliporeSigma) was added to fresh medium, and 110 µL was applied per well. Thawed lentiviral particles were equilibrated to room temperature, added to cultures, and incubated overnight at 37 °C. The following day, virus-containing medium was replaced with fresh complete medium (10% HI-FBS). Cells were rested overnight, detached with 0.25% trypsin-EDTA, neutralized, and transferred to 24-well plates for expansion. Stable transfectants were selected with puromycin (3 µg/mL, MilliporeSigma), and the medium was replaced until resistant populations were established.

### Cytokine Treatments

A549 cells were seeded in a 12-well plate at a density of 1.8 × 10^5^ cells/well and grown overnight. The next day, the cells were washed with PBS and then incubated overnight in serum-free medium (DMEM, 2% L-glutamine, and 1% Penicillin/Streptomycin). The next morning, cells were treated with a combination of 50 ng/mL TNF-α and 10 ng/mL IL-1β (PeproTech, Cranbury, New Jersey) in serum-free medium for 12 hours. RNA was then isolated using the method described in the RNA Extraction section.

### Proliferation Assay

WT, shNMC, and shPACER A549 cells, grown under standard conditions, were seeded at a density of 3.0 × 104 cells per well in 24-well plates and shaken to ensure even distribution. Methyl thiazolyl tetrazolium (MTT, Thermo Fisher), dissolved in PBS at a final concentration of 0.5 mg/mL, was added to the samples for approximately 4 hours. The MTT was removed from each well, and 500 µL of DMSO was added. Samples were covered in foil, incubated and gently shaken on a plate rocker for approximately 15 minutes. Over four days, independent wells for each timepoint were imaged on the SpectraMax iD3 (Molecular Devices) plate reader, measuring optical density (OD) at 570 nm. Empty vehicle control wells were used for background subtraction. Cells were grown in complete medium under standard conditions. Five replicates were performed per condition (n = 5).

### Migration Assay

WT, shNMC, and shPACER A549 cells were plated at 1.00 × 10^5^ cells/well in a 24-well plate and grown to near confluency. When cells were ∼90-95% confluent, the monolayer was scored with a 200 μL pipette tip to create a linear, standardized wound area. Each well was then gently washed twice with 1.5 mL PBS to remove any detached cells or debris from the wound area. Cells were maintained in low serum-containing media (1%) for the duration of the experiment. Untreated wild-type A549 cells, shNMC transduced cells and shPACER knockdown cells were imaged immediately after wound formation and at 12, 24 and 48 hours post-wound formation. The cells were imaged using a phase-contrast microscope and the Nikon NIS-Elements imaging software. Cell migration into the wound area was quantified using the "wound healing size" plugin in ImageJ (Suarez-Arnedo et al. 2020) by measuring the change in pixel area of the wound between 0 and 24 hrs.

### 3D-Spheroid Invasion Assay

Matrigel spheroids were generated by adapting a method from Aslan et al. (Aslan et al. 2021). Matrigel was thawed on a shaker at 4 °C for 1.5 hours. A549 cells (WT, shNMC, and shPACER) were detached and resuspended, then counted using a TC10 Automated Cell Counter (Bio-Rad, Hercules, CA). The cells were aliquoted into 1.5 mL tubes and centrifuged at 1000g for 5 minutes at 4 °C. The supernatant was carefully removed without disturbing the pellet, and 10 μL of Matrigel per well was added to the cell pellets. The Matrigel-cell mixture (5 × 10⁴ cells/well) was gently mixed, avoiding bubbles. The mixture was pipetted onto the center of chilled 24-well plates at a 90° angle to ensure proper spheroid formation. Plates were incubated at 37 °C for 15-20 minutes, allowing the Matrigel to solidify. After incubation, 2 mL of complete culture medium was added to the side of the plate to avoid disturbing the semi-spheres, and the plates were returned to the incubator at 37°C. The Matrigel spheroids were imaged 1 hour after plating and every 24 hours for 5 days. Top-down and cross-sectional images were collected at each point using the Nikon NIS-Elements imaging utility and software. We used these images to quantify spheroid size, invasive outgrowth, and morphological changes. Invasion was measured by analyzing the growth area relative to the initial spheroid size using the image utility ImageJ. Growth and spheroids were measured in pixel area and converted to µm or mm^2^.

### Motif Analysis Using LncLOOM

The Linux-based utility LncLOOM, created by the Ulitsky laboratory (Ross et al. 2021), was used to identify novel conserved domains within the *Homo sapiens* PACER lncRNA sequence. LncLOOMv2 was installed and run based on the instructions provided by the Ulitsky lab (GitHub repository, https://github.com/lncLOOM). The LncLOOM utility was installed on Ubuntu 20.04 LTS running on WSL2 (Windows Subsystem for Linux 2). All sequences used in our analysis were downloaded from the National Center for Biotechnology Information (NCBI) and displayed in Table 2. For each non-human primate species, a region of approximately 800bp from the antisense RNA sequence upstream of the 3′ COX-2 start site was downloaded from NCBI and aligned to the human PACER sequence. For the *Mus musculus* PACER homolog, Ptgs2os, the extended 5′ exonic region was included. The input sequences were ordered based on evolutionary distance from the target sequence prior to motif and miRNA target site predictions. LncLOOMv2 was run using the following parameters: k-mer length of 6-15, Gurobi Optimizer (Optimization, 2024), empirical statistical analysis (100 iterations), and de novo TargetScan analysis (McGeary et al. 2019). LncLOOM calculated p-values (p = 0.01) for each conserved motif based on the probability of finding that motif in a randomized version of the input sequences, where the randomized input sequences maintain the same percentage identity as the original.

### In Vivo and Ex Vivo 2A3 Treatment

A549 cells, cultured as described above, were grown to ∼75-80% confluence in a 150mm culture dish. The cells were washed with 10 mL of PBS (Gibco), resuspended in 6 mL of 0.05% Trypsin-EDTA (Thermo Fisher) and counted. For in vivo samples, 15 × 10^5^ cells were transferred to a 15 mL culture tube. The cells were pelleted by centrifugation at 400 × g for 5 minutes. The supernatant was then removed, and the cells were washed twice in PBS. The cell pellet was resuspended in 945 μL of PBS and split evenly into two 1.5 mL microcentrifuge tubes, to which 10 μL of RNase inhibitor (SUPERase-In, ThermoFisher) was added. 50 μL of 1 M, 2-aminopyridine-3-carboxylic acid imidazolide (2A3), synthesized following a previously published protocol (Incarnato 2023), was added to the modified sample, while the other sample was treated with 50 μL of DMSO as a vehicle control. The samples were then placed on a rotating shaker at 37 °C for 15 minutes. Samples were centrifuged at 16,500 × g, 4 °C for 2 minutes to pellet cells, and the supernatant was removed. RNA was extracted from the in vivo samples as previously described in the RNA Extraction section. For ex vivo samples, RNA was extracted and isolated from 7.5 × 10^5^ cells (per treatment) using the method described above; however, RNA pellets from the ex vivo samples were resuspended in 945 μL of ex vivo RNA folding buffer containing NaCl (200mM), HEPES (50mM, pH 8.0), and MgCl_2_ (5mM). Treatment with 2A3 or DMSO was performed as previously described for the in vivo samples. After 2A3 or DMSO treatment, RNA was pelleted. 10 μg of RNA from each sample was transferred to a fresh 200 μL tube and treated with 1 μL TURBO DNase (ThermoFisher) according to the manufacturer’s protocol. The DNase enzyme was inactivated with DNase inactivation reagent (ThermoFisher), followed by centrifugation at 10,000 × g for 1.5min. The DNase-treated RNA was transferred to a fresh tube, purified using the MinElute PCR Purification Kit (Qiagen) and quantified using the Qubit 4 Fluorometer (Invitrogen).

### SuperScript II Reverse Transcription

Reverse transcription of SHAPE-treated and control samples using Super Script II Reverse Transcriptase (SSII, ThermoFisher) was performed as follows: 4 μL of 500ng/μL (2μg) DNase-treated RNA, 5 μL of dH_2_O, 1 μL of 10mM dNTPs (ThermoFisher) and 1 μL of 2μM amplicon-specific primer (REV-RT, 5′-CCTGTCTGATCCCTCCCTCT-3′, synthesized by MilliporeSigma) were combined and mixed in a 200 μL PCR tube, incubated at 65 °C for 5 minutes, then 4 °C for 2 minutes. 4 μL of freshly prepared 5 × manganese Reverse transcription buffer, with a final concentration of 250 mM Tris-HCl (pH 8.3), 375 mM KCl, and 15 mM MnCl2, 2 μL of 0.1 mM DTT (ThermoFisher) and 1 μL SUPERase·In, was added and incubated at room temperature (22 °C) for 2 minutes. 2 μL of SSII was added, then incubate as follows: 42 °C for 3 hours, 70 °C for 15 minutes, and held at 12 °C until stored short-term at -20 °C or used immediately for downstream steps.

### SHAPE-MaP Library Preparation & Sequencing

PACER lncRNA was amplified using a tiled dual amplicon strategy. Amplicon 1 and amplicon 2 were amplified using Q5 Hot Start High-Fidelity DNA Polymerase (New England Biolabs) for both stages of the two-stage PCR, following the manufacturer’s instructions with modifications (Supplemental Table 4a). Each 26 μL amplicon PCR contained 8 μL cDNA template, 5× Q5 buffer, 0.5 μL dNTPs, 1.25 μL each of forward and reverse primers (10 μM), 0.25 μL Q5 Hot Start High-Fidelity polymerase, and nuclease-free water to volume. For amplicon 1 reactions, 5 μL of nuclease-free water was replaced with 5 M Ultra-Pure Betaine (Thermo Fisher). Amplicon PCR products were purified using the AMPure XP DNA Cleanup beads (Beckman Coulter) with a low 1.0 × bead-to-sample ratio to minimize adapter carryover.

Indexing PCRs were performed in 50 μL reactions containing 23.5 μL purified amplicon PCR product, 5 μL UD Index primers (Set D, Illumina), 1 μL dNTPs, 10 μL 5× Q5 buffer, 0.5 μL Q5 Hot Start High-Fidelity polymerase, and nuclease-free water to volume; for Amplicon 1, 10 μL of water was replaced with 5M betaine (ThermoFisher). Cycling parameters are listed in Supplemental Table 4b. The PCR products were purified as previously described and eluted in 20 μL of nuclease-free water. Libraries were quantified using the Qubit 1X dsDNA HS Assay Kit (ThermoFisher) on the Qubit 4 Fluorometer, and library quality was assessed using the dsDNA 910 Reagent Kit 35-1500bp (Agilent) on a 48-capillary Fragment Analyzer 5300, where percent adapter < 1% was calculated for all samples. Using Qubit concentrations, samples were normalized to 2 nM using Resuspension Buffer with Tween 20 (Illumina).

PACER libraries were pooled independently and combined with multiplexed libraries at higher concentration to offset size-related sequencing bias. The expected loss of base call quality from large-scale homogenous sequences was addressed through the introduction of a control library (25% PhiX control spike), co-sequencing with high diversity multiplexed amplicon libraries, and use of Illumina’s XLEAP-chemistry flow cell. Final sequencing reaction concentrations are as follows: 405 pM for PACER libraries, 270 pM for high diversity libraries, 225 pM for PhiX Control v3 (Illumina). Sequencing performance achieved an average Q30 of 92.52% with 78.97% of reads passing the filter. Primer sequences, including Illumina adapter sequences and binding regions, are described in Supplemental Figure 2 and Supplemental Table 2. Primer binding sites were masked prior to downstream analysis. The amplicon span corresponds to the genomic coordinates Chr1:186,680,654-186,681,409 (GRCh38.p14; NC_000001.11). Paired-end FASTQ files were generated after sequencing.

### Processing & Analysis of SHAPE-MaP data

Initial adapter trimming, quality, and length filtering were performed using fastp (Chen 2023) with the following additional parameters: cut_front, cut_tail, cut_mean_quality 25, length_required 30. Filtered sequences were aligned to the PACER reference sequence (NC_000001.11 Reference GRCh38.p14: 186680654-186681446) using the Bowtie2 Aligner (Langmead and Salzberg 2012) integrated into ShapeMapper v2.1.5 (Busan & Weeks 2018) and depth, coverage, mutational frequencies and SHAPE reactivities were calculated using ShapeMapper v2.1.5. SHAPE reactivities were normalized to transcript depth and mutational background of the DMSO controls. ShapeMapper was run with default settings in amplicon mode, with primer masking enabled. Coverage-weighted global mutation rates were calculated as, 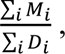 where M is the per nucleotide mutation rate and D is the depth at each position. Comparison of normalized ex vivo and in vivo conditions was performed using ΔSHAPE (Smola et al. 2015a). ΔSHAPE calculates nucleotide-wise ΔSHAPE (Ex Vivo - In vivo), applies windowed smoothing, and evaluates the significance (Z-score) of reactivity changes relative to the local background. Calculation of global paring probabilities and structural analysis was performed on both ex vivo and in vivo data sets using SuperFold (Siegfried et al. 2014) (https://github.com/Weeks-UNC/Superfold) run with default settings. SuperFold performs windowed SHAPE-directed partition-function folding across the full length of the PACER amplicon using the Deigan pseudo-free-energy model (Deigan et al. 2009). The windows were stitched together to generate global pairing probability matrices, nucleotide-resolution folding profiles, and inferred SAHPE-informed thermodynamic secondary structures.

Condition-dependent conformational heterogeneity, not characterized by global folding strategies, was analyzed using Rsample-smp, a stochastic method, and RsampleCluster (Reuter & Mathews 2010) to generate, quantify, and summarize SHAPE-restrained structural ensembles for both PACER in vivo and ex vivo conditions. Rsample-smp computed the partition function (SHAPE-weighted set of possible pairing probabilities) from the SHAPE-restrained RNA sequence, generating a partition function file (.pfs). The *Stochastic* program then generated n number of ensemble structures (n = 1000) from the probabilistic distribution encoded in the .pfs file. Finally, RsampleCluster grouped the ensemble structures using the Ding & Lawrence algorithm (Subbaramaiah et al. 2003), calculated the optimal number of clusters, and reported representative centroid structures for each group, enabling quantitative characterization of structural subpopulations within and between conditions. Jaccard index (*J*), defined as the number of shared base pairs (intersection) divided by the total number of unique base pairs across both structures (union), was calculated by comparing the base-pair sets from two structures (e.g., centroids or consensus), 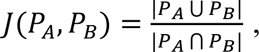 where P_A_ and P_B_ represent the sets of base pairs predicted for two distinct structures. This produced a value between 0 (no overlap) and 1 (identical), used to quantify structural similarity between predicted conformations. Base pairs were considered equivalent if |*i* − *i*′| ≤ 2 and |*j* − *j*′| ≤ 2 to allow for a ± 2 nt shift between predicted structures.

### General Statistics

All statistical analyses of cell-based assays were performed using *GraphPad Prism* (version 10.4) (GraphPad Software). All qRT-PCR and cell-based assays represent the average of three or more independent biological replicates. For qRT-PCR assays, each biological replicate was measured with n ≥ 3 technical replicates per target gene per independent experiment. The two-tailed Welch’s t-test was performed when comparing two independent conditions, while one-way ANOVA with Bonferroni corrected two-tailed Welch’s t-test (when significant) was used when measuring more than two independent conditions. For all cell-based assays, p < 0.05 is considered statistically significant. If not explicitly discussed, the statistical basis for these programs can be found in the referenced publications and repository links

## Acknowledgments

We want to thank all current and former members of the Shorrock and Lutz laboratories who participated in this work. We thank the RNA Institute Next Generation Transcriptomic Facility at the University at Albany for performing RNA sequencing.

## Author contributions

Dr. Carol S. Lutz is the corresponding author for this manuscript. Samuel Z. Desind and Samira K. Bell performed all cell culture experiments. The laboratory of Dr. Shorrock, with Robert Merritt and Benjamin Carr, contributed to the development and execution of our SHAPE-MaP protocol, sequencing approach, and analysis pipeline. All authors have intellectually contributed to the experimental work, read and approved this final manuscript for publication.

## Conflicts of Interest

Authors have no conflicts of interest to declare.

## Funding

This research was funded by the Rutgers School of Graduate Studies and the following grants: The New Jersey Commission on Cancer Research (NJCCR, COCR23PRF015), the New Jersey Health Foundation (PC161-23) and the National Institute of Health (1R01NS135254 & K99/R00NS124994 to HKS). The contents of this article are solely the responsibility of the authors and do not necessarily represent the official views of the National Institutes of Health or other funding agencies.

## Figure Legends

**Supplemental Figure 1.**
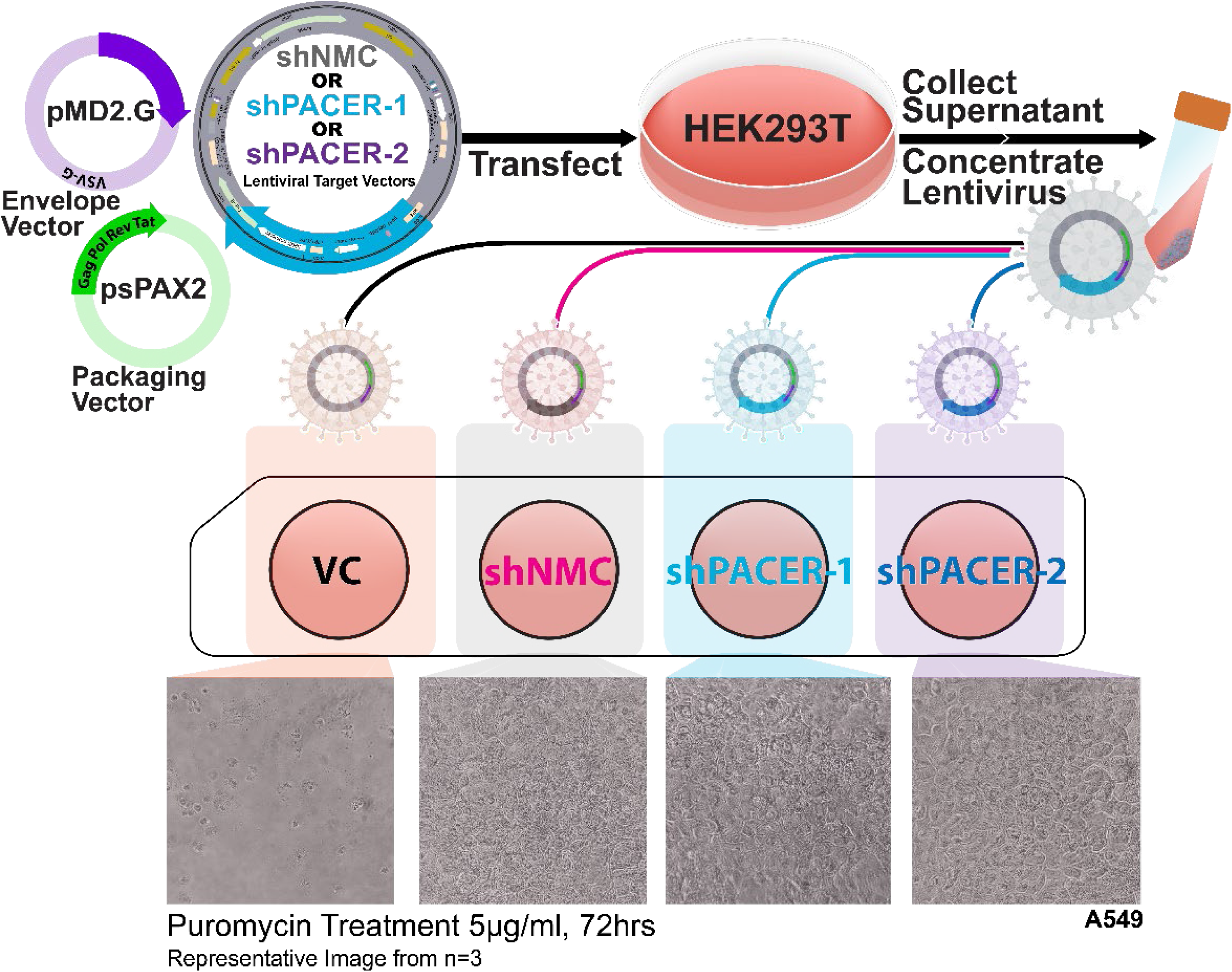
Formation of Lentivirus & Lentiviral Transduced Cell Lines. Lentiviral particles were generated by co-transfecting HEK293T cells with a target vector (non-mammalian control shRNA, shNMC; PACER-targeting shRNAs, shPACER-1, or shPACER-2), the packaging plasmid (psPAX2), and envelope plasmid (pMD2.G). Supernatants from HEK293T cells containing viral particles were collected, concentrated, and used to transduce wild-type A549 cells at multiple concentrations for each condition or PACER-targeting shRNAs (shPACER-1, shPACER-2). Cells were selected with 5 µg/mL puromycin for 72 hours alongside a vehicle control (VC). Representative images were captured with a light microscope at 100 × total magnification.

**Supplemental Table 1.**
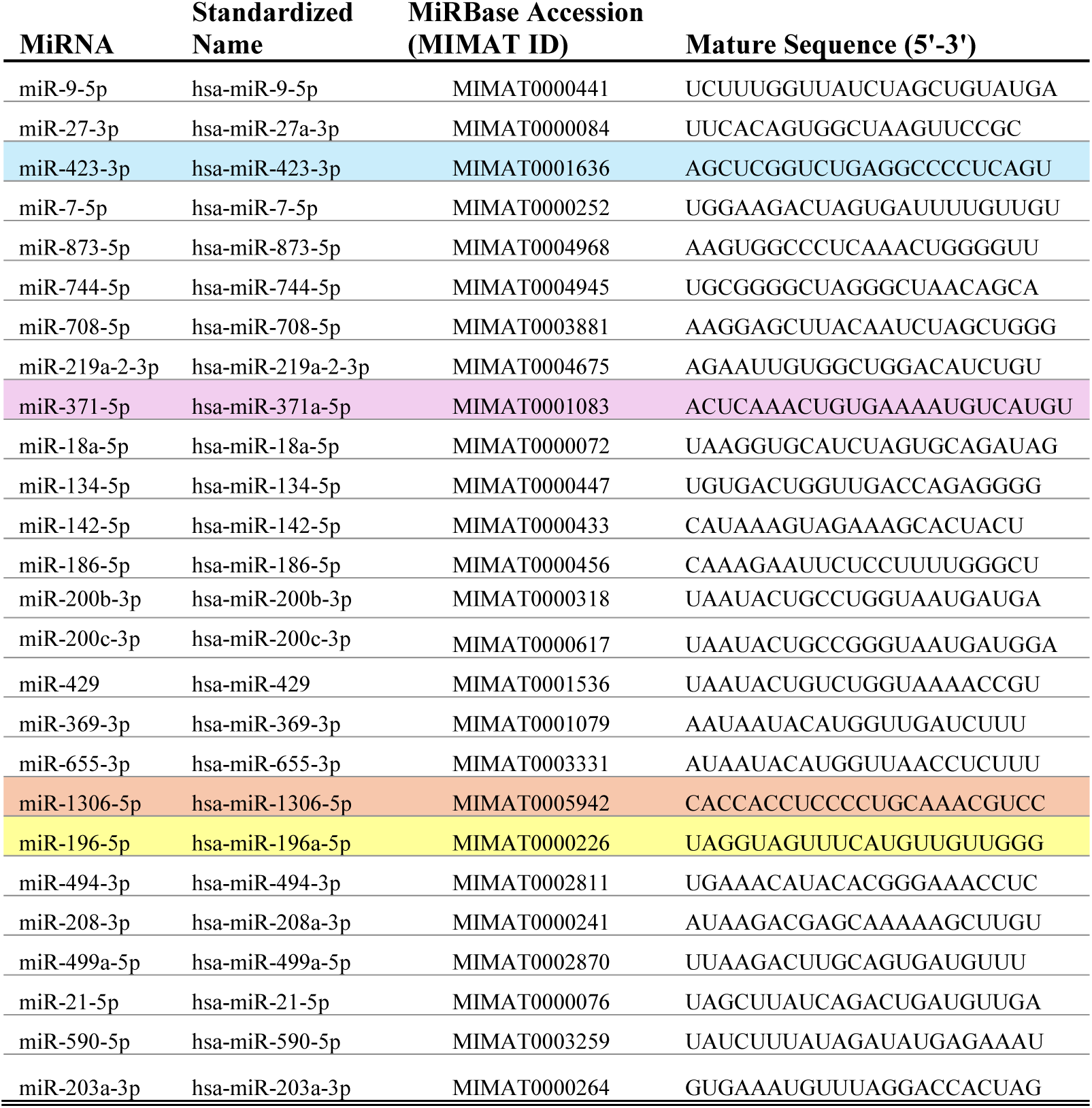
Conserved Motifs at Depth 7 With Predicted miRNA Target Sites. LncLOOM analysis and Target Scan prediction of miRNA binding sites identified 26 conserved miRNA binding sites conserved to a depth of seven. These predicted miRNA binding sites are conserved in all primate sequences but not in mice. Of these, four miRNA binding sites (highlighted, colors matching Figure 5) are conserved in all primates and conserved in mice.

**Supplemental Figure 2.**
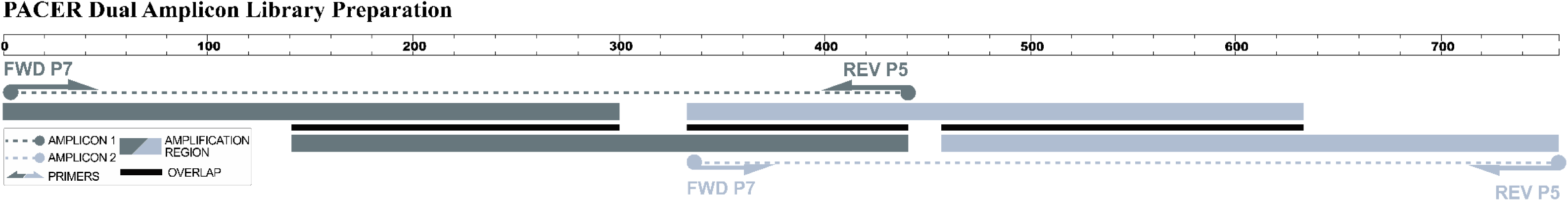
Dual Amplicon Library Preparation & Primer Tiling Strategy. Two partially overlapping amplicons tile the PACER transcript to ensure complete coverage. Amplicon 1 (dark grey) spans 1-440 nts, Amplicon 2 (light grey) spans 333-756 nts, and there is a 108 nt central overlap (nts 333-440, black bars) between the amplicons. The dotted lines show the total coverage of each amplicon, while the solid bars indicate the regions amplified by two sets of primers. Primer binding sites and direction are indicated by arrows (FWD P7 and REV P5) for each amplicon, and these positions were masked prior to downstream analysis.

**Supplemental Table 2.**
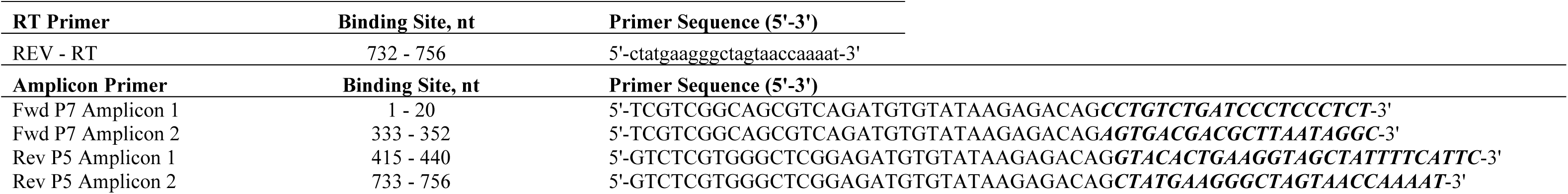
SHAPE Reverse Transcription & Amplicon Primers. Table 2 shows the complete primer sequence of primers used in primer-specific SSII reverse transcription (REV – Reverse transcription) and in the generation of PACER amplicons for SHAPE-MaP analysis. Primers are shown 5′ to 3′ and include the Illumina overhangs (in capital letters). The binding site column indicates the PACER region bound by the annealed portion of each primer. The binding region of each primer is bold and italicized. The amplicon sequence spans genomic coordinates, Chr 1:186,680,654 -186,681,409 (GRCh38.p14).

**Supplemental Figure 3.**
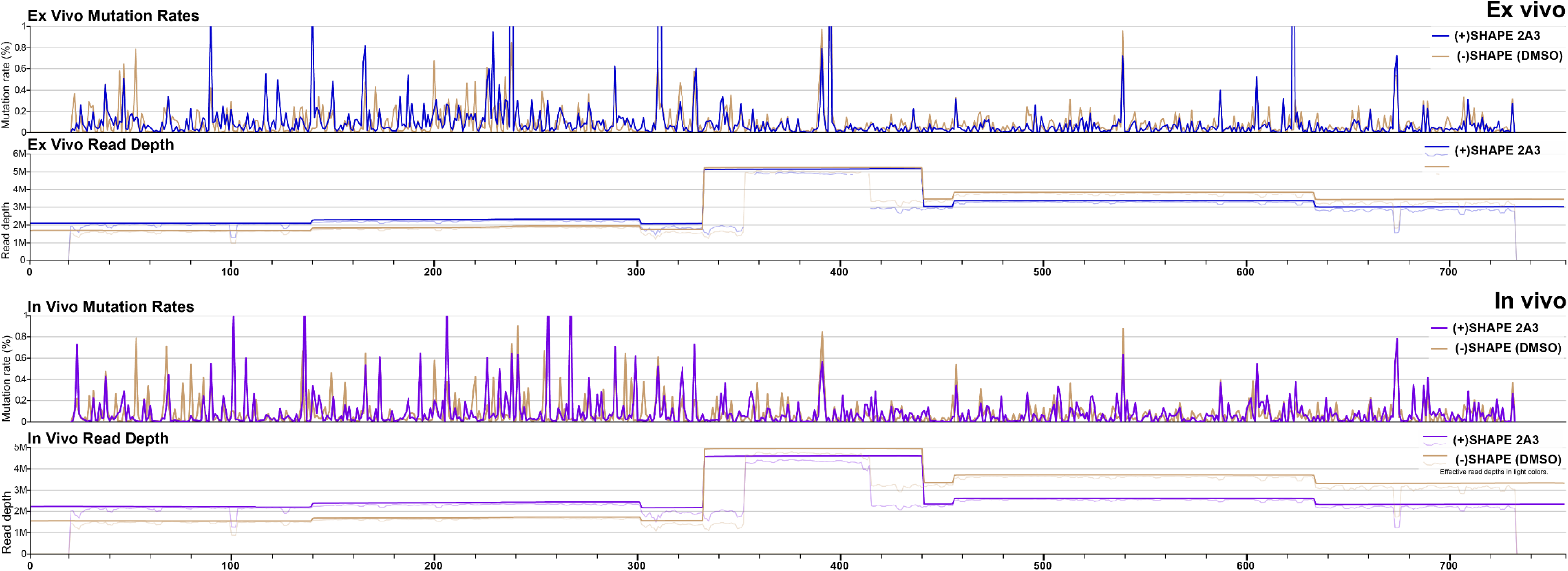
SHAPE-MaP Sample Mutation Rates & Sequencing Coverage Profiles. Per-nucleotide mutation frequencies were calculated by ShapeMapper v2.1.5 from A549 cells treated with 2A3 (+SHAPE, blue/purple) or vehicle control (−SHAPE, tan). Line plots show raw mutation rates across the PACER transcript. Coverage tracks display total mapped read depth (thick line; median depth per nucleotide: ex vivo DMSO, 1.8 × 106; ex vivo 2A3, 2.2 × 106; in vivo DMSO, 1.7 × 106; in vivo 2A3, 2.2 × 106) and effective read depth after preprocessing and alignment (lighter shaded line). All libraries achieved complete amplicon coverage and read depths well above established ShapeMapper thresholds for accurate reactivity estimation.

**Supplemental Table 3.** Shapemapper Summary Metrics. Summary of sequencing and ShapeMapper analysis metrics for ex vivo and in vivo PACER libraries treated with 2A3 (+SHAPE) or vehicle control (DMSO, −SHAPE). Global mutation rates (mean per-nucleotide mutation frequency, coverage-weighted) were elevated in 2A3-treated libraries compared to controls, confirming efficient adduct formation. The fraction of nucleotides with mutation rates above background is reported as "Mutation Rate Above Background." Alignment rates indicate the percentage of reads mapped to the PACER reference sequence. No positions exceeded thresholds for high background. The fraction of highly reactive nucleotides (>0.5 reactivity) was low, consistent with PACER’s highly structured nature. The median per-nucleotide read depth for both 2A3 and DMSO libraries was well above the minimum threshold, with 100% of nucleotides meeting the coverage sufficiency.

|  | Global Mutation Rate |  | Mutation Rate<br>Above<br>Background | Alignment Rate |  | High<br>Background | Highly Reactive<br>Nucleotides | Median Per nt<br>Read Depth |  | Depth<br>Sufficiency |
| --- | --- | --- | --- | --- | --- | --- | --- | --- | --- | --- |
|  | 2A3 | DMSO |  | 2A3 | DMSO |  |  | 2A3 | DMSO |  |
| **Ex vivo** | **0.0881%**<br>($8.81 \times 10^{-4}$) | **0.0767%**<br>($7.67 \times 10^{-4}$) | 48.30% | 92.60% | 96.30% | 0.00% | 1.10% | 2,267,704 | 1,897,934 | 100% (666/666) |
| **In vivo** | **0.0764%**<br>($7.64 \times 10^{-4}$) | **0.0707%**<br>($7.07 \times 10^{-4}$) | 54.10% | 96.40% | 92.50% | 0.00% | 1.00% | 2,299,772 | 1,761,541 | 100% (666/666) |

**Supplemental Table 4a.**
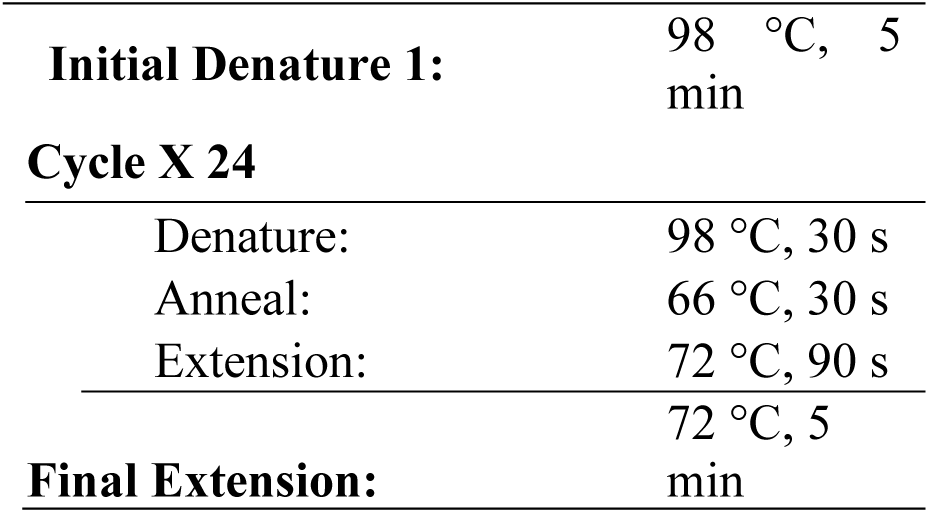
Library Amplicon PCR Protocol. PCR cycling protocol for preparation of amplicons 1 and 2 using cDNA from the SSII primer-specific reverse transcription step. Amplicon 1 was reacted with 1 M betaine to enhance amplification for both amplicon PCR and indexing PCR, while Amplicon 2 received no Betaine.

| **Supplemental Table 4a. Amplicon PCR Protocol** |  |
| --- | --- |
| **Initial Denature 1:** | 98 °C, 5 min |
| **Cycle X 24** |  |
| Denature: | 98 °C, 30 s |
| Anneal: | 66 °C, 30 s |
| Extension: | 72 °C, 90 s |
| **Final Extension:** | 72 °C, 5 min |

**Supplemental Table 4b.** Library Indexing PCR Protocol. PCR cycling protocol for indexing with the DNA/RNA UD Indexes Set D (Illumina) Tagmentation indexing primers. Amplicons 1 and 2 were run in separate reactions to limit sample-indexing bias.

| **Supplemental Table 4b. Indexing PCR Protocol** |  |
| --- | --- |
| **Initial Denature 1:** | 98 °C, 5 min |
| **Cycle X 12** |  |
| Denature: | 98 °C, 30 s |
| Anneal: | 55 °C, 30 s |
| Extension: | 72 °C, 90 s |
| **Final Extension:** | 72 °C, 5 min |

**Supplemental Table 5.**
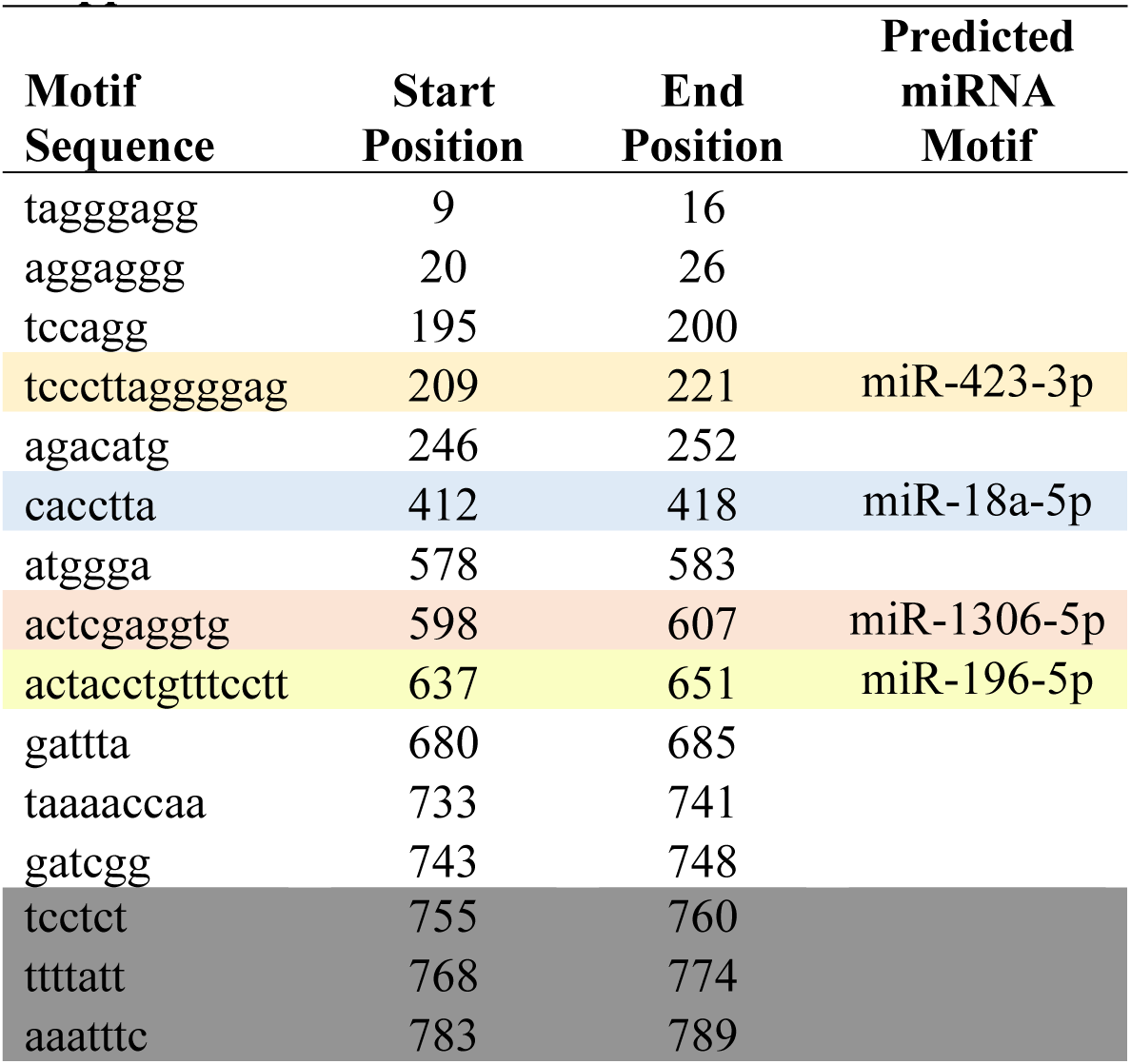
Motifs Conserved at All Levels: *Homo sapiens* - *Mus musculus*. LncLOOM analysis identified 16 motifs conserved at all levels (*Homo sapiens* - *Mus musculus*), 4 of which match predicted miRNA binding motifs. Grayed-out rows lack SHAPE- MAP reactivity and structural pairing data.

| **Motif**<br>**Sequence** | **Start**<br>**Position** | **End**<br>**Position** | **Predicted**<br>**miRNA**<br>**Motif** |
| --- | --- | --- | --- |
| tagggagg | 9 | 16 |  |
| aggaggg | 20 | 26 |  |
| tccagg | 195 | 200 |  |
| tcccttaggggag | 209 | 221 | miR-423-3p |
| agacatg | 246 | 252 |  |
| cacctta | 412 | 418 | miR-18a-5p |
| atggga | 578 | 583 |  |
| actcgaggtg | 598 | 607 | miR-1306-5p |
| actacctgttcctt | 637 | 651 | miR-196-5p |
| gattta | 680 | 685 |  |
| taaaaccaa | 733 | 741 |  |
| gatcgg | 743 | 748 |  |
| tcctct | 755 | 760 |  |
| ttttatt | 768 | 774 |  |
| aaatttc | 783 | 789 |  |

## Notes

### Competing Interest Statement

The authors have declared no competing interest.

